# Ageing compromises mouse thymus function and remodels epithelial cell differentiation

**DOI:** 10.1101/2020.03.02.973008

**Authors:** J Baran-Gale, MD Morgan, S Maio, F Dhalla, I Calvo-Asensio, ME Deadman, AE Handel, A Maynard, S Chen, F Green, RV Sit, NF Neff, S Darmanis, W Tan, AP May, JC Marioni, CP Ponting, GA Holländer

**Affiliations:** MRC Human Genetics Unit, University of Edinburgh, Edinburgh, UK; Wellcome Sanger Institute, Wellcome Genome Campus, Hinxton, UK; EMBL-EBI, Wellcome Genome Campus, Hinxton, UK; Weatherall Institute of Molecular Medicine, University of Oxford, Oxford, UK; Department of Paediatrics, University of Oxford, Oxford, UK; Cancer Research UK; Department of Biomedicine, University of Basel, and University Children’s Hospital, Basel, Switzerland; Chan Zuckerberg Biohub, San Francisco, CA; Cambridge Institute, Li Ka Shing Centre, University of Cambridge, Cambridge, UK; Department of Biosystems Science and Engineering, ETH Zurich, Basel, Switzerland

## Abstract

Ageing is characterised by cellular senescence, leading to imbalanced tissue maintenance, cell death and compromised organ function. This is first observed in the thymus, the primary lymphoid organ that generates and selects T cells. However, the molecular and cellular mechanisms underpinning these ageing processes remain unclear. Here, we show that mouse ageing leads to less efficient T cell selection, decreased self-antigen representation and increased T cell receptor repertoire diversity. Using a combination of single-cell RNA-seq and lineage-tracing, we find that progenitor cells are the principal targets of ageing, whereas the function of mature thymic epithelial cells is compromised only modestly. Specifically, an early-life precursor cell population, retained in the mouse cortex postnatally, is virtually extinguished at puberty. Concomitantly, a medullary precursor cell quiesces, thereby impairing maintenance of the medullary epithelium. Thus, ageing disrupts thymic progenitor differentiation and impairs the core immunological functions of the thymus.

## Introduction

Ageing compromises the function of vital organs via alterations of cell type composition and function (López-Otín et al., 2013). The ageing process is characterised by an upregulation of immune system associated pathways, referred to as inflamm-ageing, which is a conserved feature across tissues and species (Benayoun et al., 2019). Ageing of the immune system first manifests as a dramatic involution of the thymus. This is the primary lymphoid organ that generates and selects a stock of immunocompetent T cells displaying an antigen receptor repertoire purged of pathogenic “Self” specificities, a process known as negative selection, yet still able to react to injurious “Non-Self” antigens (Palmer, 2013). The thymus is composed of two morphological compartments that convey different functions: development of thymocytes and negative selection against self-reactive antigens are both initiated in the cortex before being completed in the medulla (Abramson and Anderson, 2017; Klein et al., 2014). Both compartments are composed of a specialized stromal microenvironment dominated by thymic epithelial cells (TECs). Negative selection is facilitated by promiscuous gene expression (PGE) in TEC, especially so in medullary TEC (mTEC) that express the autoimmune regulator, AIRE (Sansom et al., 2014). This selection ultimately leads to a diverse but self-tolerant T cell receptor (TCR) repertoire.

Thymic size is already compromised in humans by the second year of life, decreases further during puberty, and continuously declines thereafter (Kumar et al., 2018; Linton and Dorshkind, 2004; Palmer, 2013). With this reduced tissue mass, cell numbers for both lymphoid and epithelial cell compartments decline. This is paralleled by an altered cellular organization of the parenchyma, and the accumulation of fibrotic and fatty changes, culminating in the organ’s transformation into adipose tissue (Shanley et al., 2009). Over ageing, the output of naïve T cells is reduced and the peripheral lymphocyte pool displays a progressively altered TCR repertoire (Egorov et al., 2018; Thome et al., 2016). What remains unknown, however, is whether stromal cell states and subpopulations change during ageing and, if so, how these changes impact on thymic TCR selection.

To resolve the progression of thymic structural and functional decline we studied TEC using single-cell transcriptomics across the first year of mouse life. We investigated how known and previously unrecognized TEC subpopulations contribute to senescence of the stromal scaffold and correspond to alterations of thymocyte selection and maturation. Our results reveal transcriptional signatures in mature TEC subtypes that recode these cells’ functions during ageing. Unexpectedly, we discovered that the loss and quiescence of TEC progenitors are major factors underlying thymus involution. These findings have consequences for targeted thymic regeneration and the preservation of central immune tolerance.

## RESULTS

### Thymus function is progressively compromised by age

Thymus morphological changes were evident by 4 weeks of age in female C57BL/6 mice, including cortical thinning and the coalescence of medullary islands (Figure 1a). These gross tissue changes coincided with changes in thymocyte and TEC cellularity (Figure 1b,c), as noted previously (Gray et al., 2006; Manley et al., 2011). Total thymic and TEC cellularity halved between 4 and 16 weeks of age (Figure 1b,c).

**Figure 1:**
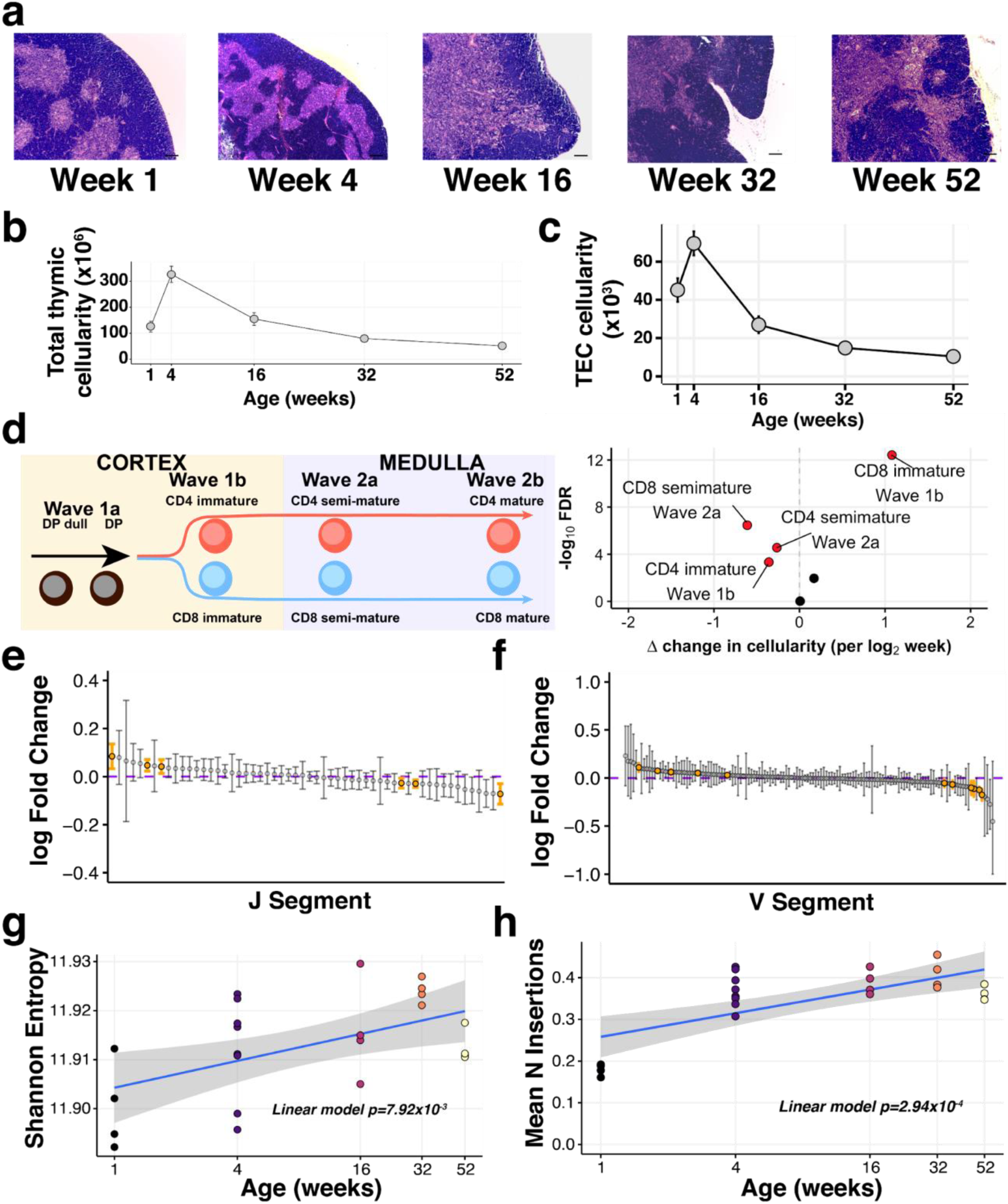
The decline of thymic cellularity and immune function with age. (a) Age-dependent changes in thymic architecture, as shown by representative H&E staining of thymic sections. Scale bars represent 150µm. Medullary islands stain as light purple while cortical regions stain as dark purple. (b) Total and (c) TEC cellularity changes in the involuting mouse thymus. Error bars represent mean +/− standard error (5 mice per age). (d) Thymocyte negative selection declines with age: (Left) Schematic showing the progression of T cell development and Wave selection stages in the thymus that were investigated; (Right) Volcano plot showing the differential abundance of each of these thymocyte negative selection populations over age. Populations that are statistically significantly altered with age (FDR 1%) are labelled and highlighted in red. (e-f) The distribution of log-fold changes showing the alterations in TCR J segment (e) and V segment (f) usage with (log_2_) age. Log fold changes +/− 99% confidence intervals are plotted, with differentially abundant segments coloured in orange. (g) Mature thymocyte TCR repertoire diversity changes with age. The y-axis indicates the Shannon entropy of M2 thymocyte TCR CDR3 clonotypes at each age (n=4-7 mice per time point), derived from TCR-sequencing of ∼15,000 cells per sample. P-value has been calculated from a linear model that regresses Shannon entropy on log age. (h) The number of non-templated nucleotide insertions detected by TCR sequencing increases with age. The displayed P-value is from a linear model that regresses mean number of inserted nucleotides on log age.

Given this sharp decline in TEC cellularity, we investigated whether the primary function of the thymus was compromised. Using flow cytometry we profiled developing thymocytes undergoing negative selection (Methods; Supplementary Figure 1a), a process that can be partitioned into four key stages: (1) double positive thymocytes (Wave 1a: Helios+PD-1+), (2) immature CD4+/CD8+ single positive (SP) thymocytes (Wave 1b: Helios+PD-1+), (3) semi-mature CD4+/CD8+ SP thymocytes (Wave 2a: Helios+) and, (4) mature CD4+/CD8+ SP thymocytes (Wave 2b: Helios+; Figure 1d, left panel) (Daley and Smith, 2013). Across these control stages the frequency of negatively selected thymocytes varied with age (Figure 1d, right panel). Specifically, negative selection of MHC class II restricted (i.e. CD4+ SP) thymocytes decreased after the first week of life (Wave 1b) concomitant with an increased removal of MHC class I restricted (CD8+ SP) cells (Figure 1d, Supplementary Figure 1b,c). In contrast, the proportions of both CD4+ and CD8+ semi-mature thymocytes undergoing negative selection in the medulla diminished with age (Figure 1d).

Impaired negative selection in the medulla undermines the production of a self-tolerant TCR repertoire. Using TCR-targeted bulk sequencing of the most mature CD4+ SP thymocytes (denoted ‘M2’) (James et al., 2018), we observed that 1 week old mice exhibited shorter CDR3 lengths and a lower proportion of non-productive TCR α and β chain sequences than older mice (Supplementary Figure 1d,e). V(D)J segment usage is altered by age, which has the potential to reshape the antigen specificity repertoire of newly generated T cells. Approximately one-third of β chain V or J segments showed an age-dependent use (38% and 29%, respectively), illustrating the robustness of TCR V(D)J usage to thymic involution and the decline in thymocyte negative selection. Diversity of the TCR repertoire amongst the most mature thymocytes, however, increased significantly over age (Figure 1g), along with the incorporation of more non-templated nucleotides (Figure 1h). The latter is inversely correlated with the post-puberty decline in the expression of thymocyte terminal deoxynucleotidyl transferase (Cherrier et al., 2002), suggesting an ageing-altered mechanism that is not intrinsic to the developing T cells. Taken together, these dynamic changes indicate that the principal immune functions of the thymus are progressively compromised with involution.

### Ageing remodels the thymic stromal epithelium

To determine whether the different TEC subpopulations were indiscriminately affected by ageing, we identified and analysed four major mouse TEC (CD45^-^EpCAM^+^) subpopulations at 5 postnatal ages using flow cytometry (Supplementary Table 1) (Gray et al., 2002; Wada et al., 2011): cortical TEC (cTEC), immature mTEC (expressing low cell surface concentrations of MHCII, designated mTEC^lo^), mature mTEC (mTEC^hi^) and terminally differentiated mTEC (i.e. mTEC^lo^ positive for desmoglein expression, Dsg3+ TEC) (Figure 2a & Supplementary Figure 2a). Following index-sorting, SMART-Seq2 single-cell RNA-sequencing, and quality control (Supplementary Figure 2b-h), we acquired 2,327 single-cell transcriptomes, evenly distributed across the 4 cytometrically-defined subpopulations and the 5 ages.

**Figure 2:**
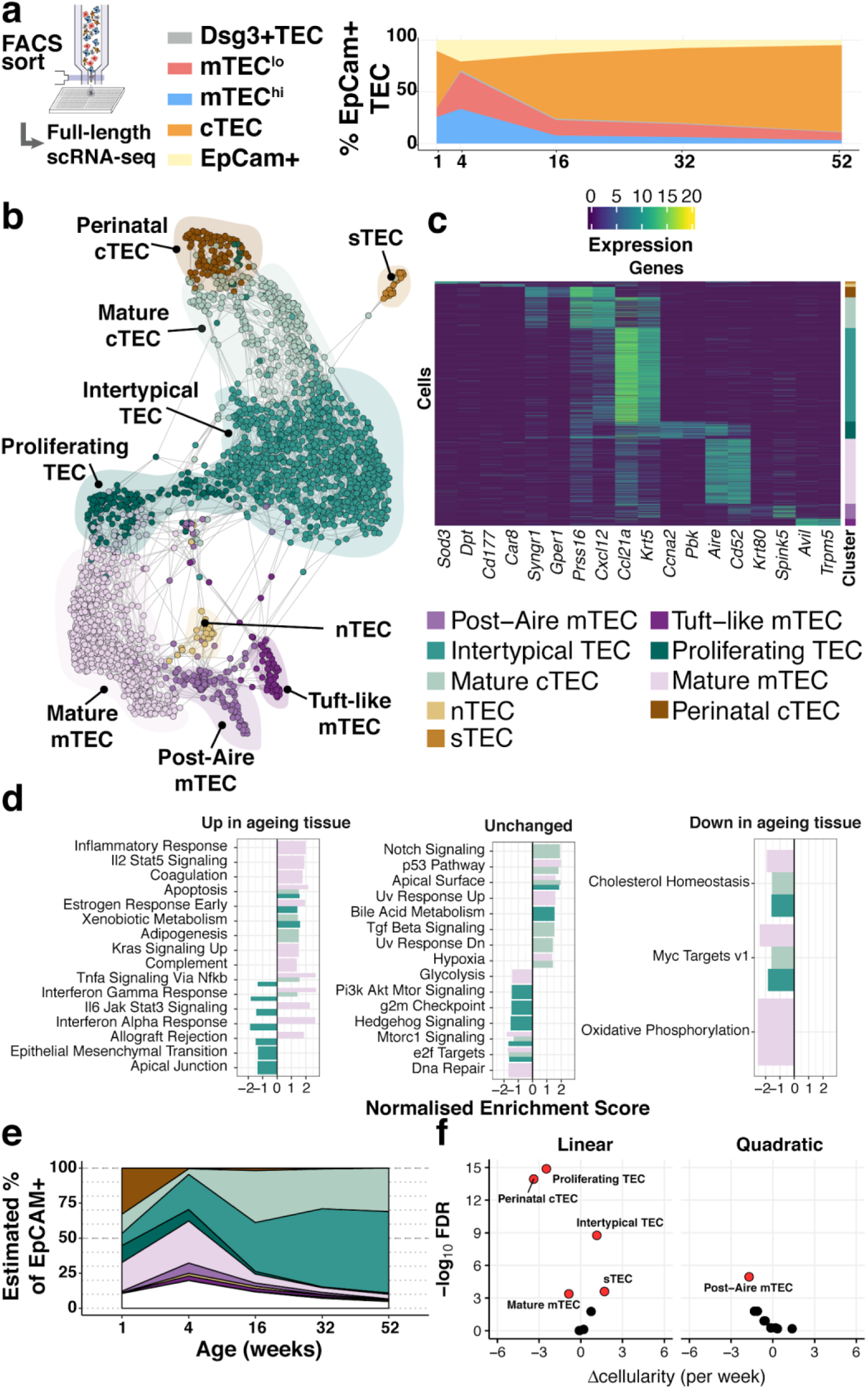
Thymic stromal remodelling during ageing. (a) A schematic showing the experimental design and FACS phenotypes of sorted cells for single-cell RNA-sequencing. Right panel shows cell composition fluctuations as a relative fraction of all EpCAM+ TEC with respect to the TEC subsets investigated. Remaining EpCAM+ cells not FAC-sorted are represented in the EpCAM+ population. (b) A SPRING-layout of the shared nearest-neighbour graph of single TEC, derived from scRNA-seq transcriptional profiles. Graph nodes represent single cells and edges represent shared k-nearest neighbours (k=5). Cells are coloured by a clustering that joins highly connected networks of cells based on a random walk (Walktrap (Pons and Latapy, 2005)). Clusters are annotated based on comparisons to known TEC subsets and stereotypical expression profiles (Table 1). (c) A heatmap of marker genes for TEC subtypes identified from single-cell transcriptome profiling annotated as in (b). (d) Enrichment of MSigDB biological pathways with age in mature cTEC, intertypical TEC and mature mTEC, annotated as in (b). Bars denote normalised enrichment score (NES) for significant pathways (FDR 5%), with enrichments coloured by cell type. Age-related alterations are shown in the context of pathways that are up-regulated (left), down-regulated (right) or do not change (middle) across multiple tissues and species (Benayoun et al., 2019). (e) A ribbon-plot demonstrating the compositional changes in TEC subtypes across ages, as an estimated fraction of all TEC (EpCAM+). Colours indicating each subtype are shown above the plot with unsorted TEC indicated in white. (f) A volcano-plot of a negative binomial generalised linear model (GLM) showing linear (left) and quadratic (right) changes in cell cluster abundance as a function of age. X-axis denotes the change (Δ) in cellularity per week, and the Y-axis shows the −log_10_ false discovery rate (FDR). Subtypes with statistical evidence of abundance changes (FDR 1%) are labelled and shown as red points.

**Table 1:** Single-cell defined TEC subtypes and known concordant phenotypes.

| TEC subtype | Locations | Populations | Reference |
| --- | --- | --- | --- |
| nTEC | Unknown<sup>1</sup> | nTEC | (Wülfing et al., 2018) |
| sTEC | Unknown<sup>1</sup> |  |  |
| Perinatal cTEC | Cortex |  |  |
| Mature cTEC | Cortex | cTEC | (Bornstein et al., 2018) |
| Intertypical TEC | Assumed:CMJ<sup>2</sup>/Cortex | mTEC I | (Bornstein et al., 2018) |
|  |  | PDPN+ CCL21+ jTEC | (Onder et al., 2015) |
|  |  | MHCII<sup>lo</sup> TPA<sup>lo</sup> TEC | (Michel et al., 2017) |
|  |  | SCA1+ PLET1+ TEC | (Ulyanchenko et al., 2016) |
|  |  | SCA1<sup>hi</sup> α6 integrin+ TEC | (Lepletier et al., 2019) |
| Proliferating TEC | Cortex & Medulla | mTEC II | (Bornstein et al., 2018) |
| Mature mTEC | Medulla | mTEC II | (Bornstein et al., 2018) |
| Post-AIRE mTEC | Medulla | mTEC III | (Bornstein et al., 2018) |
|  |  | Post-AIRE | (Nishikawa et al., 2010) |
| Tuft-like TEC | Medulla | mTEC IV | (Bornstein et al., 2018) |
|  |  |  | (Miller et al., 2018) |

<sup>1</sup>assumed cortex in adult. <sup>2</sup>Corticomedullary junction

Our analysis revealed 9 TEC subtypes (Figure 2b,c), thus providing a greater richness of epithelial states than previously reported (Bornstein et al., 2018) (Supplementary Figure 3) and a greater diversity than the 4 phenotypes cytometrically selected in this study (Supplementary Figure 4a-b). The individual subtypes were distinguished by the expression of marker genes (Figure 2c, and Supplementary Table 2), including some that are well established (post-AIRE mTEC: *Krt80, Spink5*; Mature cTEC: *Prss16, Cxcl12*; mature mTEC: *Aire*, *Cd52*) and others that have been described more recently (Tuft-like mTEC: *Avil, Trpm5*) (Bornstein et al., 2018; Miller et al., 2018). Importantly, each TEC subtype, as defined by its single-cell transcriptome (Figure 2b), did not segregate exclusively with a single cytometrically defined TEC population (Supplementary Figure 4a, Table 1). For example, a subtype that we termed intertypical TEC (*Ccl21a, Krt5*; Table 1), and which was evident at all postnatal time-points, was composed of cells from each of the four cytometrically defined TEC subpopulations. Hereafter, for clarity, we refer to transcriptomically-defined TEC clusters as subtypes and cytometrically-specified TEC as subpopulations.

Four novel TEC subtypes were identified (Table 1): perinatal cTEC (marked by the expression of *Syngr1*, *Gper1*), intertypical TEC (*Ccl21a, Krt5*) and two rare subtypes, termed neural TEC (nTEC: *Sod3*, *Dpt*) and structural TEC (sTEC, *Cd177, Car8*) based on their enrichment of neurotransmitter and extracellular matrix expression signatures (e.g. *Col1a1, Dcn, Fbn1*), respectively (Supplementary Figure 5). Specifically, nTEC both lacked expression of *Rest* (RE1 silencing transcription factor), a transcriptional repressor that is typically expressed in all non-neuronal cells (Nechiporuk et al., 2016), and expressed genes silenced by REST (*Snap25, Chga, Syp*). Perinatal cTEC were derived almost exclusively from the cytometric cTEC population and expressed β-5t (encoded by *Psmb11*), which is both a component of the cortical thymoproteosome and a marker of TEC progenitors (Mayer et al., 2016; Ohigashi et al., 2013). In addition to sharing many of the classical cTEC markers (Figure 2c; *Prss16, Cxcl12*), perinatal cTEC were characterised by a highly proliferative transcriptional signature (Supplementary Figure 5). In contrast, intertypical TEC were derived from both cortical and medullary subpopulations, and expressed gene markers associated with a progenitor-like TEC^lo^ phenotype (Table 1).

To investigate how involution affected expression changes within each subtype as well as relative changes in the abundance of individual subtypes, we identified genes that changed expression in an age-dependent manner (Figure 2d, Supplementary Figure 6) and modelled TEC subtype abundance as a function of age (Figure 2e,f). The cellular abundance of most TEC subtypes (6 of 9) varied significantly over age (Figure 2e,f; Methods). For example, perinatal cTEC represented approximately one-third of all TEC at week 1 (Figure 2e) but contributed less than 1% three weeks later. Conversely, the proportion of mature cTEC and intertypical TEC increased over time reaching ∼30% and ∼60% of all TEC, respectively, by 1 year.

Gene expression signatures that are characteristic of ageing across diverse organs and species have been reported (Benayoun et al., 2019). Many of these signatures were also evident in the transcriptomes of individual ageing TEC subtypes (Figure 2d). For example, as they aged, mature mTEC genes involved in inflammatory signalling, apoptosis and increased KRAS signalling were up-regulated, whereas genes involved in cholesterol homeostasis and oxidative phosphorylation were down-regulated (Figure 2d, left and right panels, respectively). In contrast, intertypical TEC displayed the opposite pattern (Figure 2d, left panel, dark green bars): their ageing-related decrease in cytokine signalling pathways contrasted with the stronger inflammatory signature characteristic of senescent tissues, a.k.a. inflamm-ageing (Franceschi et al., 2006). In summary, mouse thymus involution is mirrored by alterations in both TEC subtype composition and transcriptional states. The transcriptional signature of inflamm-ageing was restricted to mature cTEC and mTEC (Supplementary Figure 6) and altered subtype frequency was most striking for intertypical TEC and perinatal cTEC.

A principal function of mTEC is the promiscuous expression of genes encoding self-antigen and this was also altered across age (Supplementary Figure 7). In general, mRNA abundance of AIRE-dependent and -independent tissue restricted antigen-genes (TRAs) declined with age (Supplementary Figure 7a-f). Transcripts of eye-, pancreas- and tongue-restricted antigens displayed the most striking reduction in expression (Supplementary Figure 7g-h). Notable exceptions to this general pattern of reduced TRA expression were macrophage-associated transcripts involved in inflammatory cytokine signalling, as noted above, whose expression increased over age (Figure 2d & Supplementary Figure 7i). PGE of AIRE-controlled genes was diminished at later ages, even when *Aire* transcripts persisted, suggesting a mechanism of transcription that is reliant on factors other than AIRE abundance. Therefore, PGE in mature mTEC, and thus their capacity to represent “Self”, is increasingly compromised with age.

### Ageing compromises the differentiation of intertypical TEC into mature mTEC

Our newly described intertypical TEC subtype exhibits a transcriptional signature that includes marker genes for both mature cTEC and mature mTEC, as well as previously described mTEC progenitors (Supplementary Table 2). Furthermore, based on their position in a diffusion map between mature TEC states (Figure 3a), we hypothesised that these cells represent a TEC progenitor state. Mature mTEC are derived from progenitor cells located at the cortico-medullary junction which express β-5t (encoded by *Psmb11*) (Mayer et al., 2016; Ohigashi et al., 2013). To experimentally investigate whether intertypical and mature TEC share a common progenitor, we lineage traced the progeny of β-5t+ TEC using a triple transgenic mouse (denoted 3xtg^β5t^) with a doxycycline-inducible fluorescent reporter, ZsGreen (ZsG), under the control of the *Psmb11* promoter (Figure 3b; (Mayer et al., 2016; Ohigashi et al., 2013)). Forty-eight hours after doxycycline treatment, we isolated ZsG+ mTEC (Ly51-UEA1+CD86-) from a 1 week old mouse (Figure 3b) and profiled the traced cells using SMART-Seq2 scRNA-sequencing before comparing them with our reference atlas (Figure 3c-d). This revealed ZsG+ cells to be highly enriched for mature mTEC and intertypical TEC subtypes (Figure 3d-e) consistent with intertypical TEC and mature mTEC being derived from a common, β-5t+, progenitor, and with intertypical TEC including cells that have precursor potential to differentiate into mTEC.

**Figure 3.**
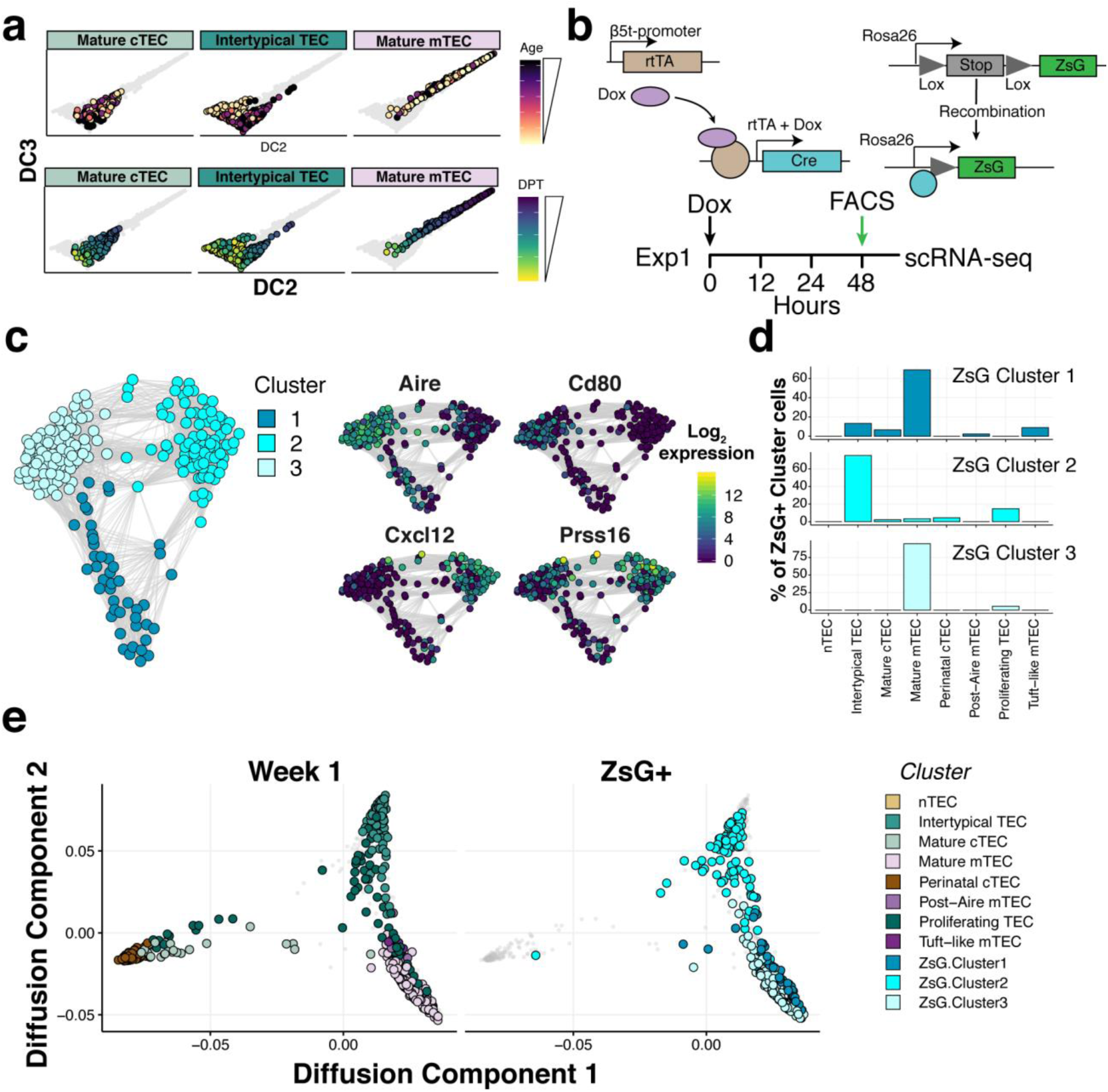
Intertypical TEC and medullar y TEC are d erived fro m a β5t+ progenitor. (a) Diffusion maps illustrating the transcriptional continuity between cortical, medullary and intertypical TEC across mouse age (top), and inferred diffusion pseudotime (DPT; bottom). (b) A schematic representing the transgenic Dox-inducible ZsGreen (ZsG) lineage tracing of β5t-expressing mTEC precursors (top), and lineage tracing experiment in 1 week old thymi (bottom). The green arrow denotes the interval post-Dox treatment. (c) A Fruchterman-Reingold layout of the SNN-graph of FAC-sorted ZsG+ mTEC from 1 week old mice, 48 hours-post Dox treatment. Graph nodes represent cells coloured by a clustering of closely connected cells. Inset panels illustrate the expression of key medullary (*Aire, Cd80*) and cortical (*Cxcl12*, *Prss16*) marker genes. (d) A β5t-expressing precursor is the common origin of intertypical TEC and mature mTEC as shown by random forest classification of ZsG+ TEC. (e) A joint diffusion map between single TEC at week 1 (left panel), and ZsG+ TEC (right panel). Points represent single cells, and are coloured by their assigned cluster as in Figure 2 (week 1 TEC) or Figure 3d (ZsG+ TEC).

Ageing intertypical TEC were characterised by progressive quiescence with age (down-regulation of Myc target genes; Figure 2d), and expression of *Itga6* (CD49f; Supplementary Table 2), a marker of quiescent, radioresistant TEC (Dumont-Lagacé et al., 2017). Consequently, we reasoned that expansion of the intertypical TEC population during ageing reflects its diminished capacity to differentiate into mature mTEC. Therefore, we used the 3xtg^β5t^ mice to explore how the relationships among progenitor, intertypical and mature mTEC change with age. TEC were labelled at weeks 1, 4 and 16 and harvested 4 weeks later in triplicate (Figure 4a & Supplementary Figures 8-10). RNA velocity analysis across single-cells collected in this experiment corroborated our conclusion that mature mTEC are derived from intertypical TEC (Figure 4a). Labelled progenitor cells in older animals were unable to differentiate fully towards mature mTEC but, instead, accumulated as intertypical TEC, consistent with a partial block during differentiation (Figure 4b, Supplementary Figure 10). By following the β-5t+ and β-5t-TEC states across age, we discovered that the gradual accumulation of intertypical TEC was specific to three of its four sub-clusters (denoted here as intertypical TEC-2, -3, or -4; Figure 4c). Of note, the intertypical TEC-3 sub-cluster is characterised by *Psmb11* expression (Figure 4d, Supplementary Figure 9), suggesting that it represents the earliest mTEC precursor state. Moreover, the intertypical TEC-2 sub-cluster specifically accumulated in the ZsG-fraction (Figure 4c) indicating either that these cells had arrested their differentiation prior to the dox-treatment and are thus more than 4 weeks old, or that they arose from a β-5t-progenitor.

**Figure 4.**
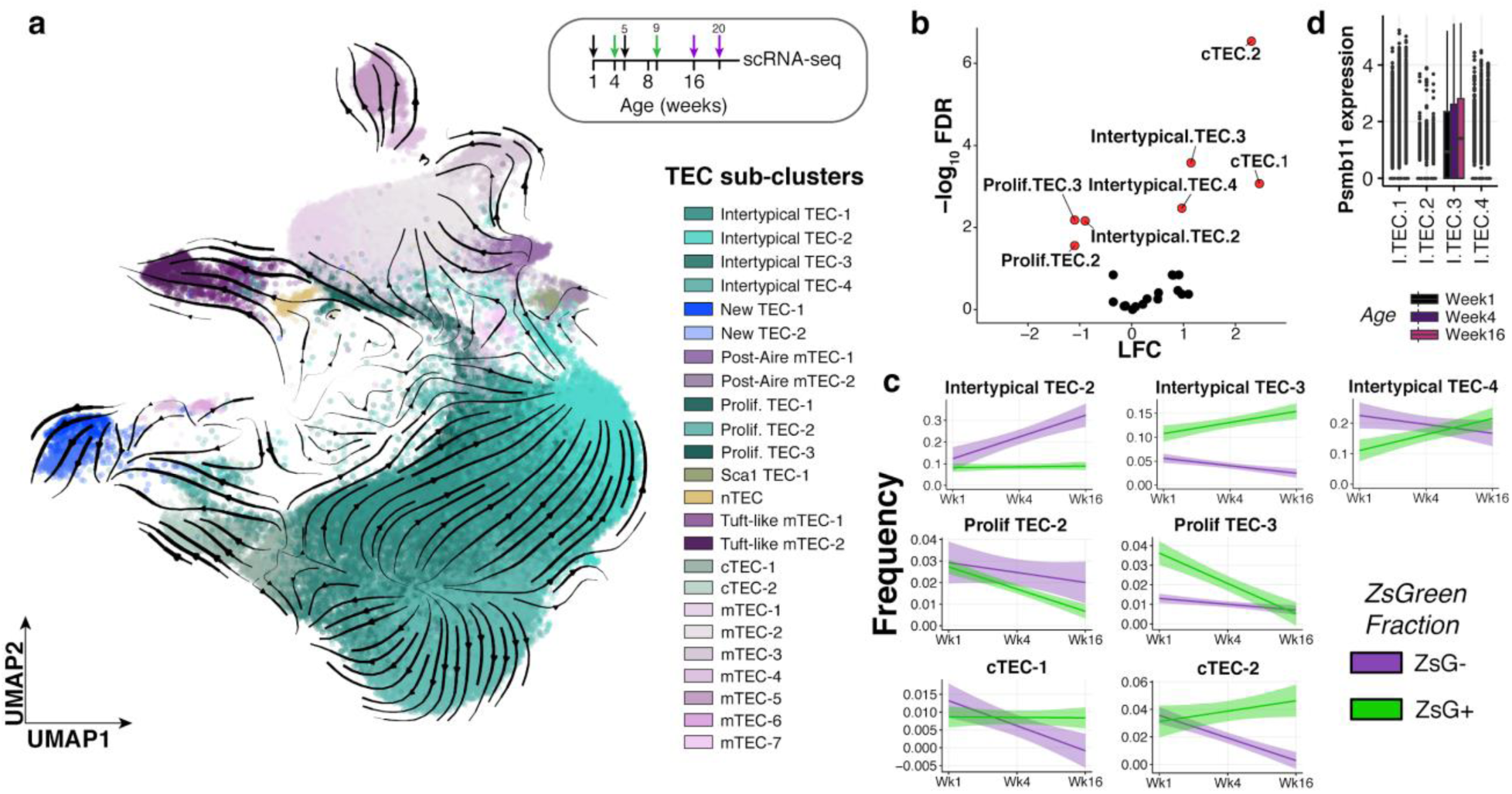
Ageing restricts the differentiation of intertypical TEC into mature mTEC. (a) RNA velocity estimates overlaid on a uniform manifold approximation and projection (UMAP) of all single cells across all ages derived from 3xtg^β5t^ mice. Cells are coloured by annotated clusters (Supplementary Figure 8) defined using a random-walk on an SNN-graph (Methods). Annotations were assigned based on the co-expression of key marker genes (Supplementary Figures 9 & 10). Inset panel: schematic representation of ZsG lineage tracing of TEC across mouse ages. Paired colour arrows denote the time and age of doxycycline treatment. Numbers above the arrows represent the age of mice at the time of single-cell measurements. (b) Differential abundance testing of TEC clusters from (a) across age and between lineage tracing fractions. The volcano plot shows the log fold change (LFC; x-axis) against −log_10_ FDR (y-axis) of the interaction between lineage fraction and age. TEC clusters that have significantly different changes in the ZsG+ compared to ZsG-fraction over age (FDR 5%) are coloured in red and labelled. Positive log-fold changes represent a higher rate of change over age in the ZsG+ fraction, whilst negative log-fold changes represent a higher rate of change in the ZsG-fraction. (c) Individual best-fit line plots show the sub-cluster frequency (y-axis) at each dox-treatment age (x-axis), grouped and coloured by ZsG fraction. The shaded band represents the linear model 95% confidence interval around the linear fit. (d) Boxplot of *Psmb11* single-cell expression (log_10_ normalised counts) across 4 intertypical TEC clusters, coloured by age at time of dox-treatment.

In summary, by combining *in vivo* lineage tracing with single-cell transcriptome profiling we have discovered that progenitor cells become increasingly blocked in intertypical TEC states during ageing and that this reduced rate of maturation results in the decline of mTEC maintenance.

## Discussion

We have demonstrated how age re-models the thymic stromal scaffold to impair its core immunological function. Leveraging the resolution of single-cell transcriptomics we identified 9 TEC subtypes, of which 4 were previously undescribed (Table 1). This refined categorization of TEC subtypes highlights the insufficiency of previously established FACS-based and ontological TEC classifications, and should facilitate detailed investigations of their function using more specific markers (Supplementary Table 2). By tracing TEC types and states across the murine life course we have found that mature TEC subtypes exhibit age-altered gene expression profiles similar to those observed across many other tissues and species (Benayoun et al., 2019). Intertypical TEC, a TEC subtype newly-defined in this study, however showed an opposing age-related pattern, with decreased expression in cytokine signalling pathways. Alongside the age-dependent decline in thymus cellularity, we observed how PGE in mature mTEC also waned over time.

Bi-potent TEC have been described with distinctive molecular identities (e.g. β-5t expression) from the postnatal thymus where they dynamically expand and contribute to the mTEC scaffold (Bleul et al., 2006; Ucar et al., 2014; Ulyanchenko et al., 2016; Wong et al., 2014). During mouse development these TEC progenitors arise from the endoderm of the third pharyngeal pouch (mid-gestation) and subsequently develop into lineage-restricted cTEC and mTEC progenitors (Baik et al., 2013; Gordon et al., 2004; Hamazaki et al., 2007; Ohigashi et al., 2013; Ripen et al., 2011; Rodewald et al., 2001; Rossi et al., 2006; Shakib et al., 2009). Using lineage tracing, we revealed how intertypical TEC arise from a β-5t+ TEC progenitor population and are a precursor to mature mTEC (Figure 3e). Thus, intertypical TEC form a previously missing link in mTEC differentiation from β-5t+ progenitors. The ability of β-5t+ TEC progenitors to expand and maintain the mTEC scaffold is progressively reduced in adolescent mice (Mayer et al., 2016). Our combined observations that intertypical TEC accumulate during ageing and up-regulate a quiescent expression signature, along with the concomitant decline in mature mTEC, are consistent with a diminished expansion and maintenance of the TEC scaffold (Figure 2d,e). Moreover, we observed that an intertypical TEC sub-cluster (intertypical TEC-3) both expresses β-5t (thus likely representing the earliest mTEC precursor) and expands with age as the population of mature mTEC contracts (Figure 4c,d). These observations indicate that the age-related expansion of this intertypical TEC sub-cluster is a direct consequence of their failure to differentiate into mature mTEC. This begs the question of what molecular mechanism leads to this defect? A recent study (Lepletier et al., 2019), suggests that TEC progenitors are re-programmed by interactions between BMP, Activin A and follistatin. In our data *Fst* (encoding follistatin), *Bmp4* and *Inhba* (encoding Activin A) are specifically expressed in the intertypical TEC compartment (Supplementary Figure 11). If the model proposed by Lepletier *et al*. is correct, then TEC progenitors may be the architects of their own malfunction.

The re-modelling of TEC maturation and the progression of inflam-ageing both alter thymus function and result in increased TCR diversity with age (Figure 1g). Concomitantly, two processes - diminution of mature TEC cellularity and blockage of TEC maturation - contribute to reduced presentation of self-antigens to developing thymocytes and thus to a less efficient negative selection. This impairment is in keeping with features of age-related thymic involution: its overall reduction in naïve T-cell output and an increased release of self-reactive T-cells (Goronzy and Weyand, 2003; Palmer, 2013). To compound these effects, the involuting thymus is also rapidly purged of its distinctive perinatal cTEC population (Figure 2e). The consequences of this are likely to be a further loss of antigen presenting cTEC and reduced support of thymocyte maturation. Taken together, we expect these TEC changes to impair the maintenance of central tolerance and could explain, at least in part, the increased incidence of autoimmunity with advancing age (Candore et al., 1997), in which the cumulative dysfunction of thymic central tolerance over time generates a slow drip feed of self-reactive T cells into the periphery.

In summary, our results reveal how the population and transcriptional dynamics of epithelial cell precursors across mouse life are coupled to age-related decline in thymic function. An enhanced understanding of the molecular mechanisms that prevent progenitors from fully progressing towards mature mTEC should facilitate studies exploring therapeutic interventions that reverse thymic decline.

## Materials and Methods

### Mice

Female C57BL/6 mice aged 1 week, 4 weeks, 16 weeks, 32 weeks, or 52 weeks were obtained from Jackson Laboratories, and rested for at least one week prior to analysis. 3xtg^β5t^ mice [β5t-rtTA::LC1-Cre::CAG-loxP-STOP-loxP-ZsGreen] mice were used for lineage-tracing experiments as previously described (Mayer et al., 2016). All mice were maintained under specific pathogen-free conditions and according to United Kingdom Home Office regulations or Swiss cantonal and federal regulations and permissions, depending where the mice were housed.

### Isolation of thymic epithelial cells and thymocytes

Thymic lobes were digested enzymatically using Liberase (Roche) and DNaseI (VWR). In order to enrich for TEC, thymic digests were subsequently depleted of CD45+ cells using a magnetic cell separator (AutoMACS, Miltenyi) before washing and preparation for flow cytometry. Thymocytes were isolated by physical disruption of thymic lobes using frosted microscope glass slides.

### Flow cytometry and cell sorting

Cells were stained at a concentration of 5-10 x10^6^ per 100µl in FACS buffer (2% fetal calf serum in PBS or 5% bovine serum albumin in PBS). Supplementary Table 3 provides details of antibody staining panels. Staining for cell surface markers was performed for 20 minutes at 4°C, except for CCR7 which was performed for 30 minutes at 37°C in a water bath prior to the addition of other cell surface stains. The FoxP3 Transcription Factor Staining Buffer Kit (eBioscience) was used according to the manufacturer’s instructions in order to stain for intracellular antigens. Cell viability was assessed using DAPI staining or LIVE/DEAD Fixable Aqua Dead Cell Stain (Invitrogen). Samples were acquired and sorted using a FACS Aria III (BD Biosciences). For single-cell RNA-sequencing index sorting was used and cells were sorted into 384 well plates. Flow cytometry data was analysed using FlowJo V 10.5.3.

### TCR rearrangement simulations

Simulations of TCR germline rearrangements were used to estimate TCR-sequencing sample sizes. Sequential steps of α- and β-chain rearrangement were simulated to model β-selection and double negative thymocyte maturation prior to negative selection. We uniformly sampled V(D)J segments from the C57BL/6 TCR locus. For the TCR β-chain, variable (V) and diversity (D) segments were randomly selected from available sequences. For joining (J) segments, the TRBJ1 locus was selected on the first attempt, and TRBJ2 if a second attempt to rearrange was made. Consequently the matching TRBC segment was selected based on the J segment that was chosen (either TRBC1 or TRBC2). For the concatenation of each segment pair, i.e. V-J, V-D, VD-J, randomly selected nucleotides were inserted between the adjoining segments, based on sampling from a Poisson distribution with λ=4. The productivity of the rearranged β-chain was determined by the presence of a complete open reading frame (ORF) beginning with a canonical start codon (‘ATG’) in the selected V segment that spanned the full V(D)J and constant segments. In the event of a failed rearrangement a second attempt was made using the TRBJ2 and TRBC2 segments. If either of these attempts produced a valid TCR β-chain, then under the principle of allelic exclusion the simulation proceeded to the α-chain rearrangement. However, if the second rearrangement failed to produce a valid TCR β-chain, the process was repeated for the second allele.

For the TCR α-chain, variable (V) and joining (J) regions were randomly selected from the available TCRA sequences. Following the same principle as above, if the simulated rearrangement failed to generate a valid TCR with a complete ORF spanning the V segment to the constant region then the simulation switched to the second allele. A successful TCR germline was recorded only in the event of both valid α- and β-chains. The complete simulation resulted in a valid α-chain in 40.2% of simulations, and a valid β-chain in 63.1% of simulations. To calculate sample sizes for our TCR-sequencing experiments we simulated 1 million “thymocytes”, and sub-sampled 10, 100, 500, 1000, 5000, 10000, 20000, 50000 and 100000 cells, defined by a productive pair of TCR chains. To simulate replicates we ran these simulations with 10 different random initiations. To establish the required sample sizes we calculated the proportions of V(D)J segment frequencies for α- and β-chains. Additionally, we calculated the TCR diversity at each sample size using the Shannon entropy across α- and β-chain CDR3 clonotypes, defined by the unique amino acid sequence. Results of simulations are shown in Supplementary Figure 12.

### TCR sequencing

15,000 M2 thymocytes (TCRb^hi^, CCR7+, MHCI+, CD69-, CD8-, CD4+, CD25-) were sorted and RNA extracted using the Qiagen RNeasy Micro kit. 10ng of RNA was used to prepare bulk TCR-seq libraries using the SMARTer Mouse TCR a/b Profiling Kit (Takara) according to instructions. Libraries were sequenced on a MiSeq (300 base paired-end reads). Reads were trimmed using Trimmomatic, down-sampled to the smallest library size and aligned using MiXCR (version 3.0).

### Haematoxylin and eosin (H&E) staining of thymic sections

Thymic lobes were harvested and cleaned under a dissecting microscope before being fixed in 10% Formalin (Sigma) for 12-36 hours, depending on size, and dehydrated in ethanol. After fixation the tissues were embedded in paraffin using an automated system (Tissue-Tek Embedding Centre, Sakura) and sectioned to a thickness of 8µm. H&E staining was performed using an automated slide stainer (Tissue-Tek DRS 2000, Sakura) and slides were visualised under a light microscope DM750 (Leica).

### Plate-based single-cell RNA-sequencing

#### Lysis plates

Single thymic epithelial cells were index FAC-sorted into 384-well lysis plates. Lysis plates were created by dispensing 0.4 μl lysis buffer (0.5 U Recombinant RNase Inhibitor (Takara Bio, 2313B), 0.0625% Triton X-100 (Sigma, 93443-100ML), 3.125 mM dNTP mix (Thermo Fisher, R0193), 3.125 μM Oligo-dT 30 VN (IDT, 5’AAGCAGTGGTATCAACGCAGAGTACT 30 VN-3’) and 1:600,000 ERCC RNA spike-in mix (Thermo Fisher, 4456740) into 384-well hard-shell PCR plates (Biorad HSP3901) using a Tempest liquid handler (Formulatrix). All plates were then spun down for 1 minute at 3220g and snap frozen on dry ice. Plates were stored at −80°C until used for sorting.

#### cDNA synthesis and library preparation

cDNA synthesis was performed using the Smart-seq2 protocol (Picelli et al., 2014). Briefly, 384-well plates containing single-cell lysates were thawed on ice followed by first strand synthesis. 0.6 μl of reaction mix (16.7 U/μl SMARTScribe TM Reverse Transcriptase (Takara Bio, 639538), 1.67 U/μl Recombinant RNase Inhibitor (Takara Bio, 2313B), 1.67X First-Strand Buffer (Takara Bio, 639538), 1.67 μM TSO (Exiqon, 5’-AAGCAGTGGTATCAACGCAGACTACATrGrG+G-3’), 8.33 mM DTT (Bioworld, 40420001-1), 1.67 M Betaine (Sigma, B0300-5VL), and 10 mM MgCl 2 (Sigma, M1028-10X1ML)) were added to each well using a Tempest liquid handler. Bulk wells received twice the amount of RT mix (1.2 μl). Reverse transcription was carried out by incubating wells on a ProFlex 2×384 thermal-cycler (Thermo Fisher) at 42°C for 90 min and stopped by heating at 70°C for 5 min.

Subsequently, 1.6 μl of PCR mix (1.67X KAPA HiFi HotStart ReadyMix (Kapa Biosystems, KK2602), 0.17 μM IS PCR primer (IDT, 5’-AAGCAGTGGTATCAACGCAGAGT-3’), and 0.038U/μl Lambda Exonuclease (NEB, M0262L)) was added to each well with a Tempest liquid handler (Formulatrix). Bulk wells received twice the amount of PCR mix (3.2 μl). Second strand synthesis was performed on a ProFlex 2×384 thermal-cycler using the following program: 1. 37°C for 30 minutes, 2. 95°C for 3 minutes, 3. 23 cycles of 98°C for 20 seconds, 67°C for 15 seconds, and 72°C for 4 minutes, and 4. 72°C for 5 minutes. The amplified product was diluted with a ratio of 1 part cDNA to 9 parts 10mM Tris-HCl (Thermo Fisher, 15568025), and concentrations were measured with a dye-fluorescence assay (Quant-iT dsDNA High Sensitivity kit; Thermo Fisher, Q33120) on a SpectraMax i3x microplate reader (Molecular Devices). These wells were reformatted to a new 384-well plate at a concentration of 0.3 ng/μl and a final volume of 0.4 μl using an Echo 550 acoustic liquid dispenser (Labcyte). If the cell concentration was below 0.3 ng/μl, 0.4 μl of sample was transferred. Illumina sequencing libraries were prepared using the Nextera XT Library Sample Preparation kit (Illumina, FC-131-1096) (Darmanis et al., 2017; Tabula Muris Consortium et al., 2018). Each well was mixed with 0.8 μl Nextera tagmentation DNA buffer (Illumina) and 0.4 μl Tn5 enzyme (Illumina), then tagmented at 55°C for 10 min. The reaction was stopped by adding 0.4 μl “Neutralize Tagment Buffer” (Illumina) and spinning at room temperature in a centrifuge at 3220 X g for 5 min. Indexing PCR reactions were performed by adding 0.4 μl of 5 μM i5 indexing primer, 0.4 μl of 5 μM i7 indexing primer, and 1.2 μl of Nextera NPM mix (Illumina). PCR amplification was carried out on a ProFlex 2×384 thermal cycler using the following program: 1. 72°C for 3 minutes, 2. 95°C for 30 seconds, 3. 12 cycles of 95°C for 10 seconds, 55°C for 30 seconds, and 72°C for 1 minute, and 4. 72°C for 5 minutes.

#### Library pooling, quality control, and sequencing

Following library preparation, wells of each library plate were pooled using a Mosquito liquid handler (TTP Labtech). Row A of the thymus plates, which contained bulk cells, was pooled separately. Pooling was followed by two purifications using 0.7x AMPure beads (Fisher, A63881). Library quality was assessed using capillary electrophoresis on a Fragment Analyzer (AATI), and libraries were quantified by qPCR (Kapa Biosystems, KK4923) on a CFX96 Touch Real-Time PCR Detection System (Biorad). Plate pools were normalized to 2 nM and sequenced on the NovaSeq 6000 Sequencing System (Illumina) using 2×100bp paired-end reads with an S4 300 cycle kit (Illumina, 20012866). Row A thymus pools were normalized to 2 nM and sequenced separately on the NextSeq 500 Sequencing System (Illumina) using 2×75bp paired-end reads with a High Output 150 cycle kit (Illumina, FC-404-2002).

### Single-cell RNA-sequencing processing, quality control and normalisation

Paired-end reads were trimmed to a minimum length of 75nt using trimmomatic with a 4nt sliding window with a quality threshold of 15. Leading and trailing sequences were removed with a base quality score < 3 (Bolger et al., 2014). Contaminating adaptors were removed from reads with a single seed mismatch, a palindrome clip threshold of 30 and a simple clip threshold of 10. Trimmed and proper-paired reads were aligned to mm10 concatenated with the ERCC92 FASTA sequences (Thermo Fisher Scientific) using STAR v2.5.3a (Dobin et al., 2013) and a splice-junction database constructed from the mm10 Ensembl v95 annotation with a 99nt overhang. Paired-end reads were aligned with the parameters: *--outSAMtype* BAM SortedByCoordinate *--outSAMattributes* All *--outSAMunmapped* Within KeepPairs; all other parameters used default values. Following alignment each single-cell BAM file was positionally de-duplicated using PicardTools *MarkDuplicates* with parameters: *REMOVE_DUPLICATES =* true, *DUPLICATE_SCORING_STRATEGY = TOTAL_MAPPED_REFERENCE_LENGTH* [http://broadinstitute.github.io/picard].

De-duplicated single-cell transcriptomes were quantified against exon sequences of the mm10 Ensembl v95 using featureCounts (Liao et al., 2014). Poor quality single-cell transcriptomes were removed based on several criteria: contribution of ERCC92 to total transcriptome > 40%, sequencing depth < 1×10^5^ paired-reads and sparsity (% zeros) > 97%. From this initial round of quality control 2780 cells were retained for normalisation and downstream analyses. Deconvolution-estimated size factors were used to normalise for sequencing depth across single cells, prior to a log10 transformation with the addition of a pseudocount (+1), implemented in *scran (Lun et al., 2016)*.

### Single-cell clustering and visualisation

TEC from all ages and sort-types were clustered together using a graph-based algorithm that joins highly connected networks of TEC based on the similarity of their expression profile. To enhance the differences in the expression profile of individual TEC libraries, we first applied a text frequency-inverse document frequency (TF-IDF) transform (Manning et al., 2008) to the gene-by-cell expression matrix. This transform enhances the signal from rarely expressed genes (of particular importance would be those that are promiscuously expressed in TEC), while also lessening the contribution from widely expressed genes. The transformed matrix represents the product of the gene-frequency and the inverse-cell-frequency. To compute this transformed matrix, we first assigned the gene-frequency matrix as the log2 of normalised gene-by-cell expression matrix (G_f_ = log_2_ (C); C is the normalised count matrix). Next, we computed the inverse-cell-frequency as the inverse frequency of detection of each gene (ICF_x_ = log10 (N / (1+E_x_)); N is the number of cells, E_x_ is the number of cells expressing gene *X*). Finally, the product of the gene-frequency matrix and inverse-cell-frequency was computed (GF_ICF = G_f_ * ICF). The highly variable genes from this transformed matrix were used to compute a shared nearest neighbor (SNN) graph (k=10), and the clusters were identified using a random walk (Walktrap (Pons and Latapy, 2005)) of the SNN graph. To assess the robustness of the clusters, we also clustered cells without the TF-IDF transform and using a series of alternate parameters. We computed a consensus matrix to determine how often the identified TEC subtypes co-clustered. We found that the identified TEC sub-types were robustly co-clustered regardless of the parameters of the clustering that was applied (Supplementary Figure 13). Visualisation of the connected graph was computed using the SPRING algorithm to generate a force-directed layout of the K-nearest-neighbor graph (k=5) (Weinreb et al., 2018).

### Treatment with Doxycycline

One-week old 3xtg^β5t^ mice were treated with a single i.p. injection of 0.004mg of Doxycycline (Sigma) diluted in Hank’s Balanced Salt Solution (Life Technologies), whereas older mice (four-week and sixteen-week old) were treated with two i.p. injections of Doxycycline (2mg, each) on two consecutive days during which they were also exposed to drinking water supplemented with the drug (2 mg/mL in sucrose (5% w/v)).

### Droplet-based single-cell RNA sequencing

#### Preparation of TEC suspensions for single-cell RNA-sequencing

Single thymic epithelial cell suspensions were obtained by enzymatic digestion using Liberase (Roche), Papain (Sigma) and DNase (Sigma) in PBS as described in (Kim and Serwold, 2019; Mayer et al., 2016). Prior to FAC-sorting, TEC were enriched for EpCAM-positivity using a magnetic cell separator (AutoMACS, Miltenyi), as described above. Enriched cells were then stained for the indicated cell surface antigens (Supplementary Table 3) in conjunction with TotalSeq-A oligonucleotide-conjugated antibodies (BioLegend) to allow for barcoding and pooling of different TEC subpopulations and subsequently sorted into 4 subpopulations: ZsGreen+ cTEC, ZsGreen-cTEC, ZsGreen+ mTEC, and ZsGreen-mTEC (Supplementary Figure 8a). After sorting, the cell viability and concentration of each of the cell samples collected were measured using a Nexcelom Bioscience Cellometer K2 Fluorescent Viability Cell Counter (Nexcelom Bioscience).

#### Droplet-based single-cell RNA-sequencing

Equal cell numbers were pooled from each of the samples, and a total of 30000 cells were loaded per well onto a Chromium Single Cell B Chip (10X Genomics) coupled with the Chromium Single Cell 3ʹ GEM, Library & Gel Bead Kit v3 and Chromium i7 Multiplex Kit (10X Genomics) for library preparation, according to the manufacturer’s instructions. In short, the cell suspension was mixed with the GEM Retrotranscription Master Mix and loaded onto well number 1 on the Chromium Chip B (10x Genomics). Wells 2 and 3 were loaded with the appropriate volumes of gel beads and partitioning oil, respectively, after which the Chromium Controller (10X Genomics) was used to generate nanoliter-scale Gel Beads-in-emulsion (GEMs) containing the single cells to be analysed. The fact that cell samples containing 6 different hashtag antibodies were pooled together allowed us to overload the 10X wells with 30000 cells per well, aiming for a recovery of approximately 12000 single cells (40%) per well. This also allowed us to overcome the resulting increase in doublet rate by subsequently eliminating from further analysis any cell barcode containing more than one single hashtag sequence. Incubation of the GEM suspension resulted in the simultaneous production of barcoded full-length cDNA from poly-adenylated mRNA as well as barcoded DNA from the cell surface protein-bound TotalSeqA antibodies inside each individual GEM. Fragmentation of the GEMs allowed for the recovery and clean-up of the pooled fractions using silane magnetic beads. Recovered DNA was then amplified, and cDNA products were separated from the Antibody-Derived Tags (ADT) and Hashtag oligonucleotides (HTO) by size selection. The amplified full-length cDNA generated from polyadenylated mRNA were fragmented enzymatically and size selection was used to optimise amplicon size for the generation of 3’ libraries. Library construction was achieved by adding P5, P7, a sample index, and TruSeq Read 2 (read 2 primer sequence) via End Repair, A-tailing, Adaptor Ligation, and PCR. Separately, ADT and HTO library generation was achieved through the addition of P5, P7, a sample index, and TruSeq Read 2 (read 2 primer sequence) by PCR. Sequences of the primers designed for this purpose can be found in Supplementary Tables 4 and 5.

#### Library pooling, quality control, and sequencing

Library quality was assessed using capillary electrophoresis on a Fragment Analyzer (AATI). The concentration of each library was measured using a Qubit dsDNA HS Assay Kit (ThermoFisher Scientific), and this information was then used to dilute each library to a 2nM final concentration. Finally, the different libraries corresponding to each sample set were pooled as follows: 85% cDNA + 10% ADT + 5% HTO, after which pooled libraries were sequenced on an Illumina NovaSeq 6000 using the NovaSeq 6000 S2 Reagent Kit (100 cycles) (Illumina).

### Droplet-based single-cell RNA sequencing processing, de-multiplexing and quality control

Multiplexed 10X scRNA-seq libraries were aligned, deduplicated and quantified using Cellranger v3.1.0. Gene expression matrices of genes versus cells were generated separately for each sample (i.e. each 10X Chromium chip well), as well as those for hashtag oligo (HTO) and antibody (ADT) libraries. Cells were called using emptyDrops, with a background UMI threshold of 100 (Lun et al., 2019). Experimental samples, i.e. replicates and ZsGreen-fractions, were demultiplexed using the assigned HTO for the respective sample (Stoeckius et al., 2018). Specifically, within each sample, the HTO fragment counts were normalised across cell barcodes for all relevant HTOs using counts per million (CPM). These CPMs were used to cluster cell barcodes using k-means with the expected number of singlet clusters, i.e. unique HTOs in the respective sample. To estimate a background null distribution for each HTO within a sample, we then selected the k-means partition with the highest average CPM for the HTO and excluded these cells, along with the top 0.5% of cells with the highest counts for the respective HTO. We then fitted a negative binomial distribution to the HTO counts for the remaining cells to estimate a threshold (*q*) at the 99th quantile. All cell barcodes with counts ≥ *q* were assigned this HTO. This procedure was repeated for each HTO within a sample. Cell barcodes that were assigned to a single HTO were called as ‘Singlets’, whilst cell barcodes assigned to > 1 HTO were called as ‘Multiplets’. Finally, cell barcodes with insufficient coverage across HTOs were called as ‘Dropouts’ (Supplementary Figure 8b). Only ‘Singlets’ were retained for normalisation and downstream analyses.

Poor quality cells barcodes were removed based on high mitochondrial content, defined within each sample as twice the median absolute deviation from the median mitochondrial fraction. Cell barcodes with low coverage (< 1000 UMIs detected) were also removed prior to normalisation. Finally, deconvolution-estimated size factors were calculated to normalise across single cells, then log10 transformed with a pseudocount (+1), as implemented in *scran (Lun et al., 2016)*.

### Droplet single-cell RNA sequencing clustering and annotation

Highly variable genes (HVGs) were defined across droplet single cells based on the estimated fit across cells between the mean log normalised counts and variance, at an FDR of 1×10^-7^ (Brennecke et al., 2013). The first 20 principal components (PCs) across HVGs were calculated, and used as input to construct an SNN-graph (k=31) across all single cells. These were then clustered into closely connected communities using the Walktrap algorithm (Pons and Latapy, 2005). Clusters were annotated based on the co-expression of TEC subtype marker genes (Supplementary Figures 8 & 9). Droplet single cells were visualised in reduced dimension with the first 20 PCs as input using uniform manifold approximation and projection (UMAP) (McInnes et al., 2018), with k=31 nearest neighbours and a minimum distance=0.3.

### RNA velocity

RNA velocity estimates the future state of single-cells based on a mechanistic model of transcription to identify groups of genes that are actively up-regulated, or down-regulated, based on the ratio of splice/unspliced sequencing reads (La Manno et al., 2018). We calculated the velocity of each single cell across single-droplet RNA-sequencing experiments using the stochastic model implemented in *scvelo* (Bergen et al., 2019). Velocity vectors were overlaid on a UMAP representation constructed using *scanpy* (Wolf et al., 2018).

### Diffusion map and pseudotime inference

Diffusion maps and diffusion pseudotime trajectories were constructed using a matrix of log-transformed size-factor normalized gene expression values across single cells as input, with highly variable genes to define the diffusion components, implemented in the Bioconductor package *destiny* (Angerer et al., 2016; Haghverdi et al., 2016). Diffusion maps for both ageing TEC and embryonic TEC used k=20. The ZsGreen experimental cells used k=21 and the first 20 principal components as the input to the diffusion map estimation. Diffusion pseudotime distances were computed from an index cell defined in each analysis.

### Cell type classification - MARS-seq from Bornstein *et al*

The MARS-seq counts matrix from (Bornstein et al., 2018) were downloaded from Gene Expression Omnibus (GSE103967), and normalised using deconvolution size factors (Lun et al., 2016), after removing cells with low sequencing coverage (<1000 UMIs, sparsity ≥98%). HVGs were detected, as described above, across single cells from all WT mice, the embryonic time points E14.5 & E18.5, 6 day old WT mouse, as well as the *Aire* and *Pou2f* knock-out mice at an FDR 0.1%. The log-transformed normalised counts for these HVGs were used as input to PCA, and the first 10 PCs were used to construct an SNN-graph (k=20). Clusters of single cells were defined based on a random walk on this graph (Pons and Latapy, 2005). A total of 9 clusters were detected.

To map these cells across we constructed a kNN classifier (k=5) implemented in the R package *FNN*, trained on the ageing single-cell data. We first took the set of commonly expressed genes between our study and those of Bornstein *et al*., and performed a per-cell cosine normalisation on each data set. These data were used as input to classify each single cell into an ageing cluster (Supplementary Figure 3d).

### Age-dependent cluster abundance modelling

Numbers of each TEC subtype or cluster were counted per replicate and at each age. Cell counts were modelled using a linear negative binomial model, with the total number of cells captured per replicate as a model weight, implemented in the Bioconductor package *edgeR* (McCarthy et al., 2012; Robinson et al., 2010). For the ZsGreen experiment we down-sampled the counts matrix to rebalance the ZsGreen+ and ZsGreen-fractions to equal proportions. We then tested the hypothesis that the interaction between age and ZsGreen fraction was different from 1. This amounts to comparing the gradients of the two regression slopes in ZsGreen+ versus ZsGreen-cells across ages. Statistically significant age-dependent changes were tested in these models using an empirical Bayes quasi-likelihood F-test (Chen et al., 2016).

### Tissue and tissue restricted antigen gene definition

Tissue restricted antigen (TRA) genes were defined based on the specificity of their expression across a broad range of mouse tissues using the FANTOM5 cap analysis of gene expression with sequencing (CAGE-seq) data that are publicly available (http://fantom.gsc.riken.jp/data/). Tissue samples were grouped into 27 broad groups based on the annotation data (Supplementary Table 6). For each protein-coding gene (based on Ensembl identifier), the per-tissue expression level was defined as the maximum run length encoding (RLE) normalised expression level. For genes with multiple transcriptional start sites, the mean RLE expression across isoforms was first taken. The specificity of tissue expression for each gene across tissues (n) was then calculated using the tau-index (τ) (Yanai et al., 2005):

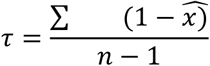

where 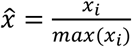.

Genes with τ ≥ 0.8 were defined as TRAs, whilst those with τ ≤ 0.4 were defined as constitutively expressed; all remaining genes were given the classification ‘miscellaneous’. Each TRA gene was assigned to one tissue, the one in which it was maximally expressed. *Aire*-dependent and -independent genes were defined using the classification from Sansom *et al*. (Sansom et al., 2014).

### Age-dependent tissue-representation modelling

The age-dependence of tissue-representation across single mTEC was tested using a negative binomial linear model. Specifically for each single mTEC the number of TRA genes with log expression > 0 was counted within each assigned tissue (see above). These single-cell tissue counts were aggregated across single mTEC at each time point, and for each replicate mouse, to yield ‘tissue counts’. Aggregated ‘tissue counts’ were then used as the dependent variable in a negative binomial linear model implemented in the Bioconductor package *edgeR*. Statistically significant age-dependent changes were defined at an FDR of 1%.

### Differential gene expression testing

All differential gene expression testing was performed in a linear model framework, implemented in the Bioconductor package *limma*. To test for age-dependent gene expression changes, log-normalized gene expression values for each gene was regressed on log2(age) and adjusted for sequencing depth for each single cell using deconvolution size factors estimated using *scran*.

### Gene signature and functional enrichment testing

Marker genes or differentially expressed genes (throughout ageing) were tested to identify enriched pathways, specifically those from MSigDB hallmark genesets or Reactome pathways. Marker genes were identified as those genes with a 4-fold enrichment in the subtype relative to all other subtypes (adjusted p < 0.01). MSigDB hallmark (Liberzon et al., 2015; Subramanian et al., 2005) and Reactome pathway (Fabregat et al., 2018) enrichments for markers of each subtype were computed using the clusterProfiler package (Yu et al., 2012). For age-dependent differentially expressed genes, gene set enrichment analysis was used (GSEA) to identify enriched MSigDB hallmark genesets. These results were categorised based on the expected change in expression due to ageing across multiple tissues and species (Benayoun et al., 2019).

### Age-dependent modelling of thymocyte negative selection

Age-dependent variation in thymocyte negative selection was modelled using a negative binomial GLM implemented in the Bioconductor package *edgeR*. Cell counts were regressed on age, using the input parent population for each replicate as a model offset to control for variation in the preceding selected population. Across populations, multiple testing was accounted for using the false discovery rate procedure (Benjamini and Hochberg, 1995), where a statistically significant relationship with age was set at 1%.

## Code and data availability

All code used to process data and perform analyses are available from https://github.com/WTSA-Homunculus/Ageing2019. All sequence data, counts matrices and meta-data are available from ArrayExpress with accession numbers E-MTAB-8560 (ageing thymus) and E-MTAB-8737 (lineage traced thymus). TCR sequencing data is available from SRA (PRJNA551022).

## Acknowledgements

We gratefully acknowledge the Chan Zuckerberg Biohub for support and for sequencing, and members of the Tabula Muris Consortium for technical assistance. CPP is funded by the MRC (MC_UU_00007/15). GH, JB and MDM were supported by the Wellcome Trust (grant 105045/Z/14/Z). JCM was supported by core funding from the European Molecular Biology Laboratory and from Cancer Research UK (award number 17197). GH and ICA was supported by the Swiss National Science Foundation (grant numbers IZLJZ3_171050 and 310030_184672). AEH was supported by a Clinical Lectureship from the NIHR. FD was supported by the Wellcome Trust [109032/Z/15/Z].

## Competing interests

CP is a reviewing editor at eLife. The authors have no further competing interests to declare.

---

**Supplementary Figure 1:**
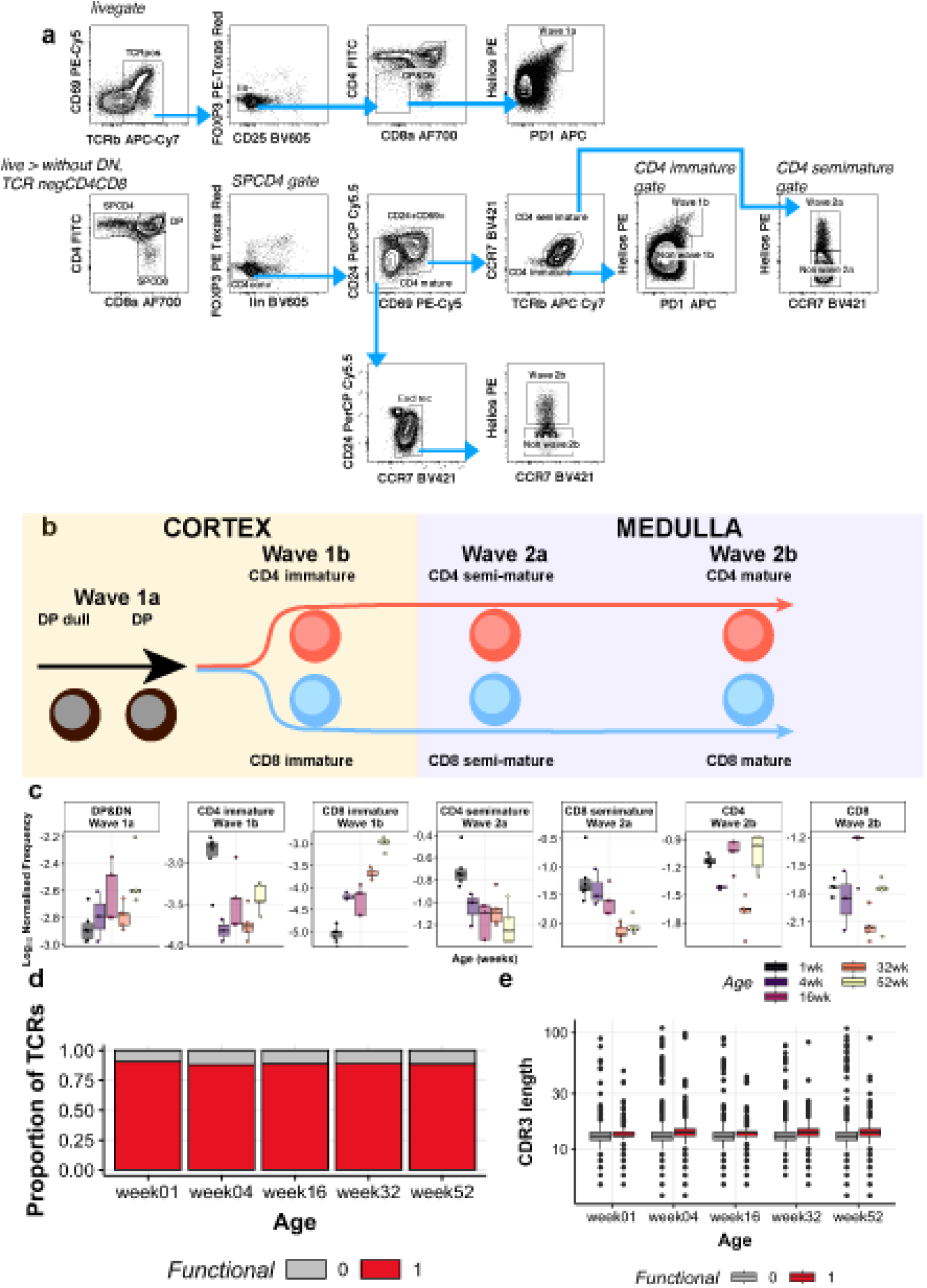
Changes in T-cell population s throughout ageing. (a) FACS gating strategy to separate different T-cell subtypes. (b) Maturation trajectory for T-cells. (c) Frequency of different subtypes by age. (d) Proportion s of viable TCRs amongst all constructe d CDR3 sequences. Functional sequences are coloured in red and non-functional sequences, defined by an incomplete sequence, premature stop codon or missing start codon, are coloured in grey. (e) CDR3 amino acid length distributions per age (X-axis). Boxplots are coloured by either functional (red) or non-functional (grey) TCR sequences.

**Supplementary Figure 2:**
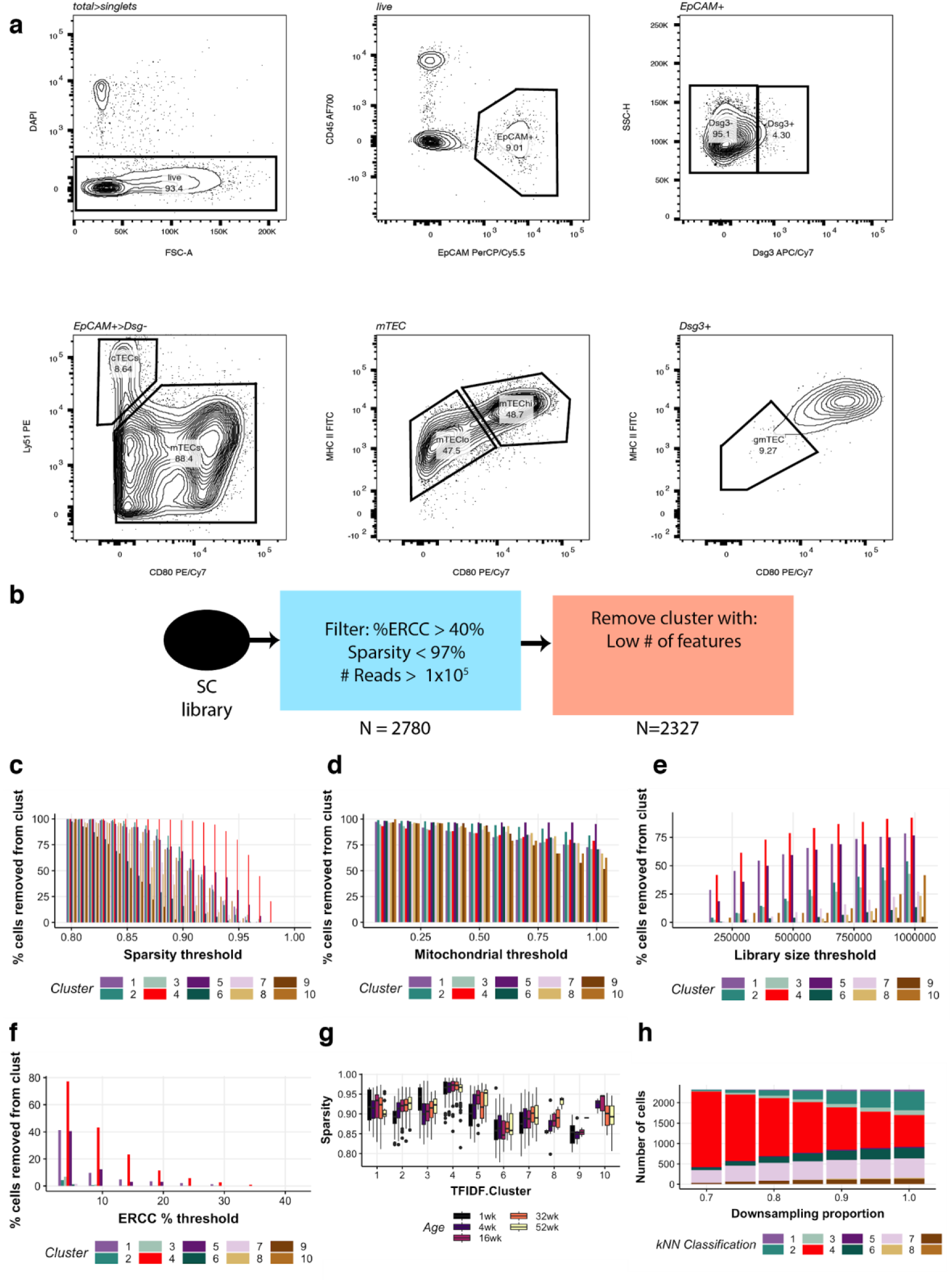
Experimental investigation of the ageing thymus. (a) FACS gating strategy for isolation of TEC sort types. (b) Filtering strategy to identify high-quality TEC libraries. (c-f) Fractions of libraries filtered out based on sparsity threshold (c), the fraction of reads from mitochondrial genes expressed (d), library size thresholds (e), or ERCC-spike in RNA % expression threshold (f). (g) Sparsity in each single cell cluster by age. (h) Reassignment of libraries to clusters based on downsampling fraction. Excessive downsampling leads to an accumulation of the low-diversity library.

**Supplementary Figure 3:**
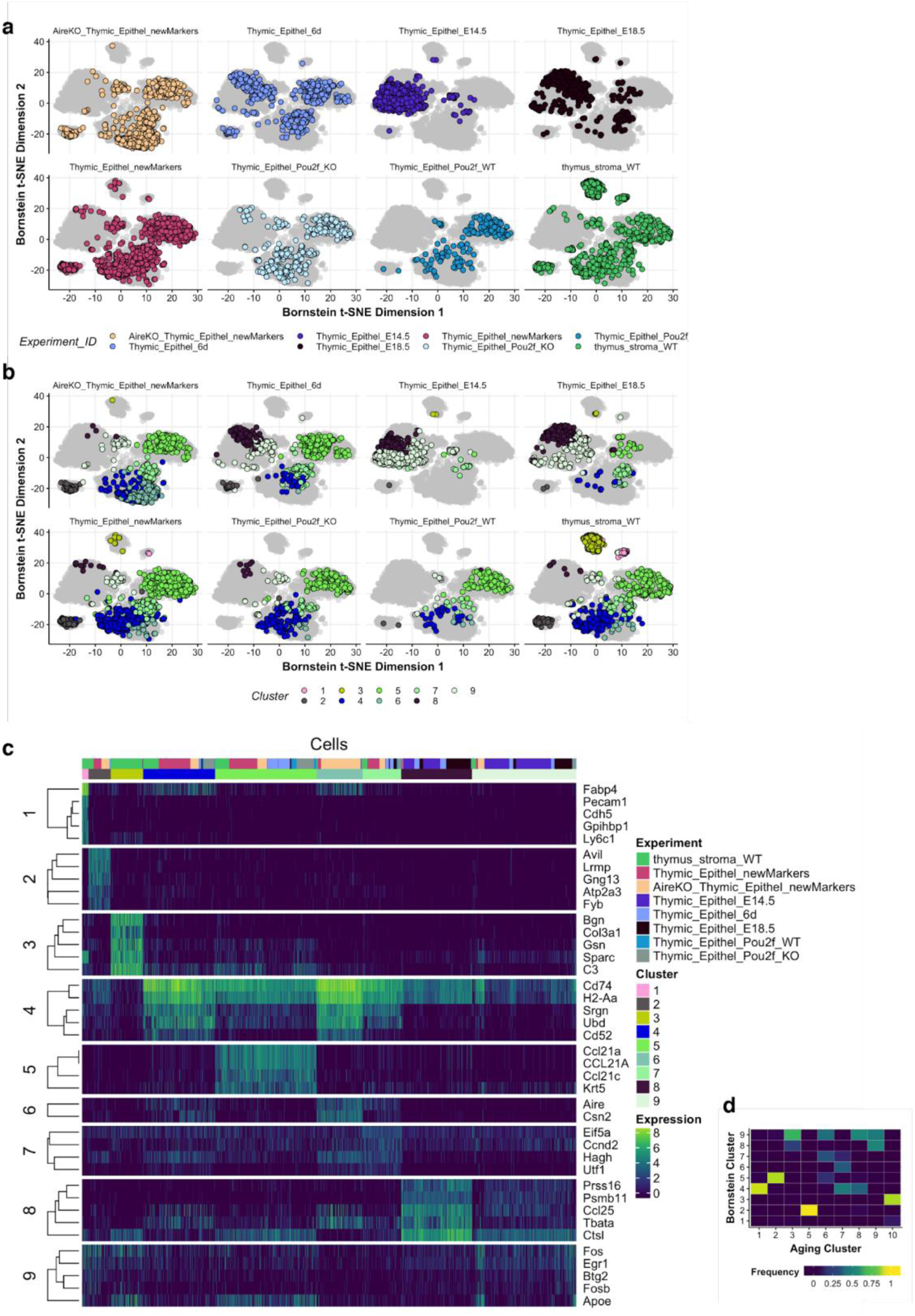
Comparison of Bornstein et al. (Bornstein et al. 2018) single-cell transcriptomes to the TEC subtypes defined in this study. (a) tSNE representation of Bornstein et al. single-cell TEC libraries. (b) Nine clusters identified from Bornstein et al. single-cell data overlaid on tSNE visualisation. (c) Expression heatmap of marker genes acquired from Bornstein et al. clusters. (d) Comparison of Bornstein et al. single-cell clusters to ageing subtypes from this study.

**Supplementary Figure 4:**
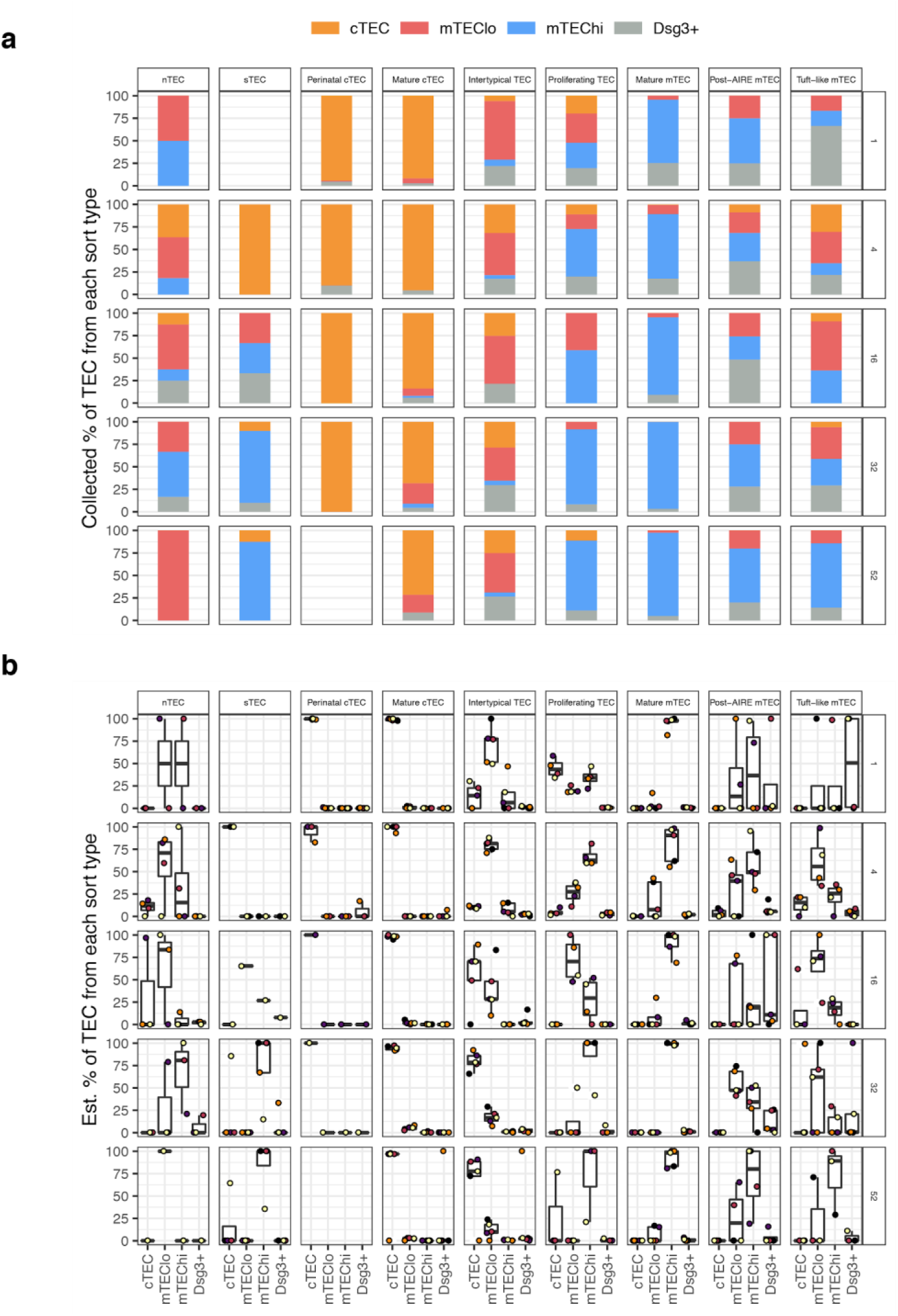
Relationship between classical FAC sort-types and transcriptionally-defined single-cell subtypes. (a) Observed percentages (%) of TEC based on pre-scoring into classical sort-types. (b) Estimated contributions of each FAC sort type to each single cell subtype through age. Each coloured dot represents data from an independent experiment.

**Supplementary figure 5:**
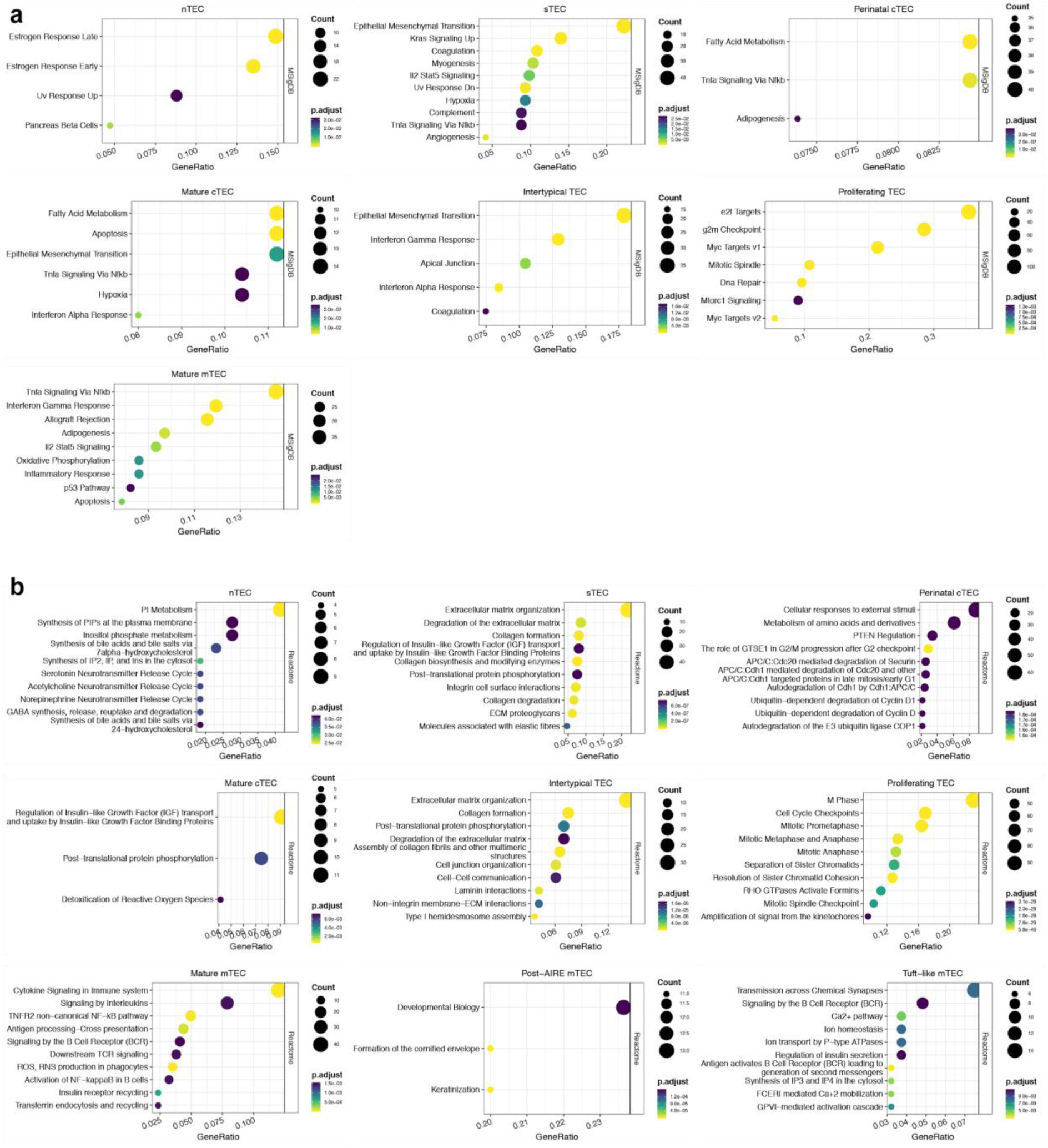
MSigDB (a) and Reactome (b) pathways enriched for expression of marker genes for each single cell subtype. The X-axis shows the fraction of marker genes that overlap the specified pathway, the size of the dot represents the number of marker genes in the enriched pathway, and the colour of the dot represents the p-value adjusted for multiple tests.

**Supplementary Figure 6:**
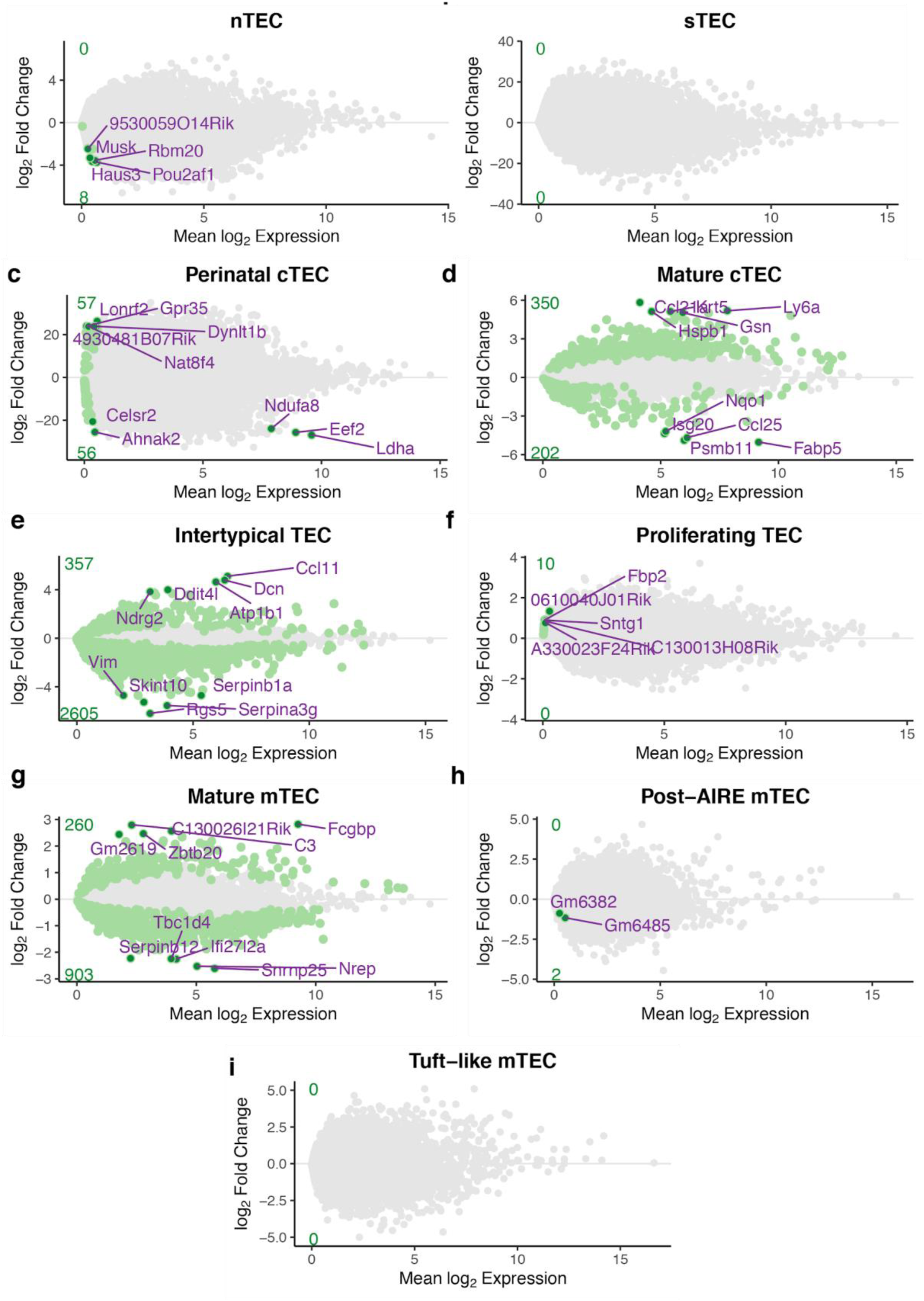
Differential expression of genes throughout ageing. Each panel shows the average expression and the log2 fold-change with age for each single cell subtype. Significantly altered genes are shown in green and the total number of up- or down-regulated genes per subtype are shown in the green font along the y-axis. The top 5 up- or down-regulated genes are labelled, where present.

**Supplementary Figure 7:**
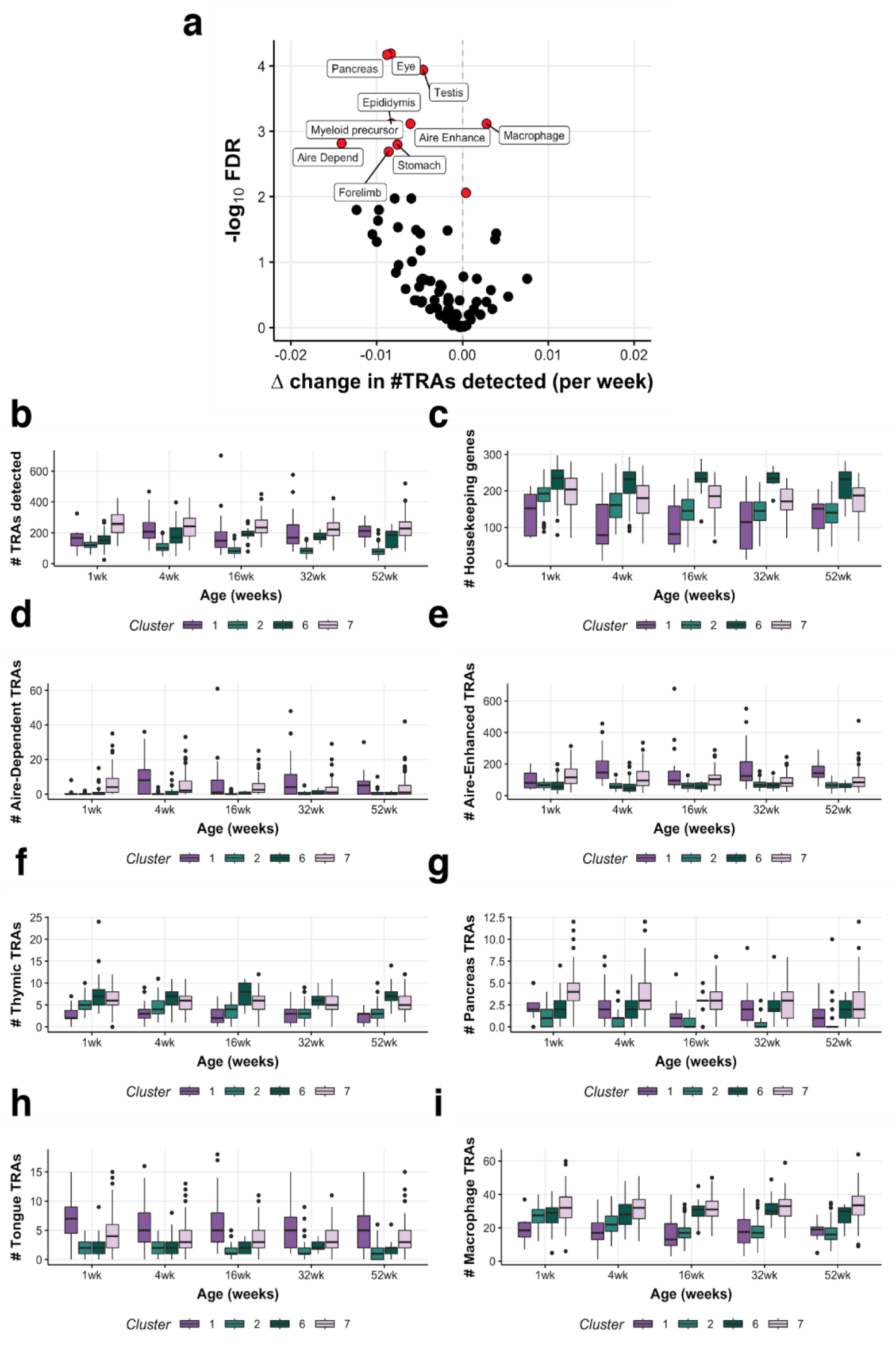
Details of tissue-restricted antigen (TRA) expression throughout ageing. (a) A volcano plot of differential TRA abundance testing, showing the consistent down-regulation of TRAs in mature mTEC. (b-i) Data for clusters 1 (Post-AIRE), 2 (Intertypical TEC), 6 (proliferating TEC), 7 (Mature mTEC) are shown as differently-coloured boxes in boxplots. The number (#) of TRAs (b), of housekeeping genes (c), of Aire-dependent TRAs (d) and of Aire-enhanced TRAs (e) expressed plotted against single-cell subtype and age of the mouse. The number of TRA genes detected from thymic (f), pancreas (g), tongue (h) and macrophage (i) specific groups by age and single-cell subtype.

**Supplementary Figure 8:**
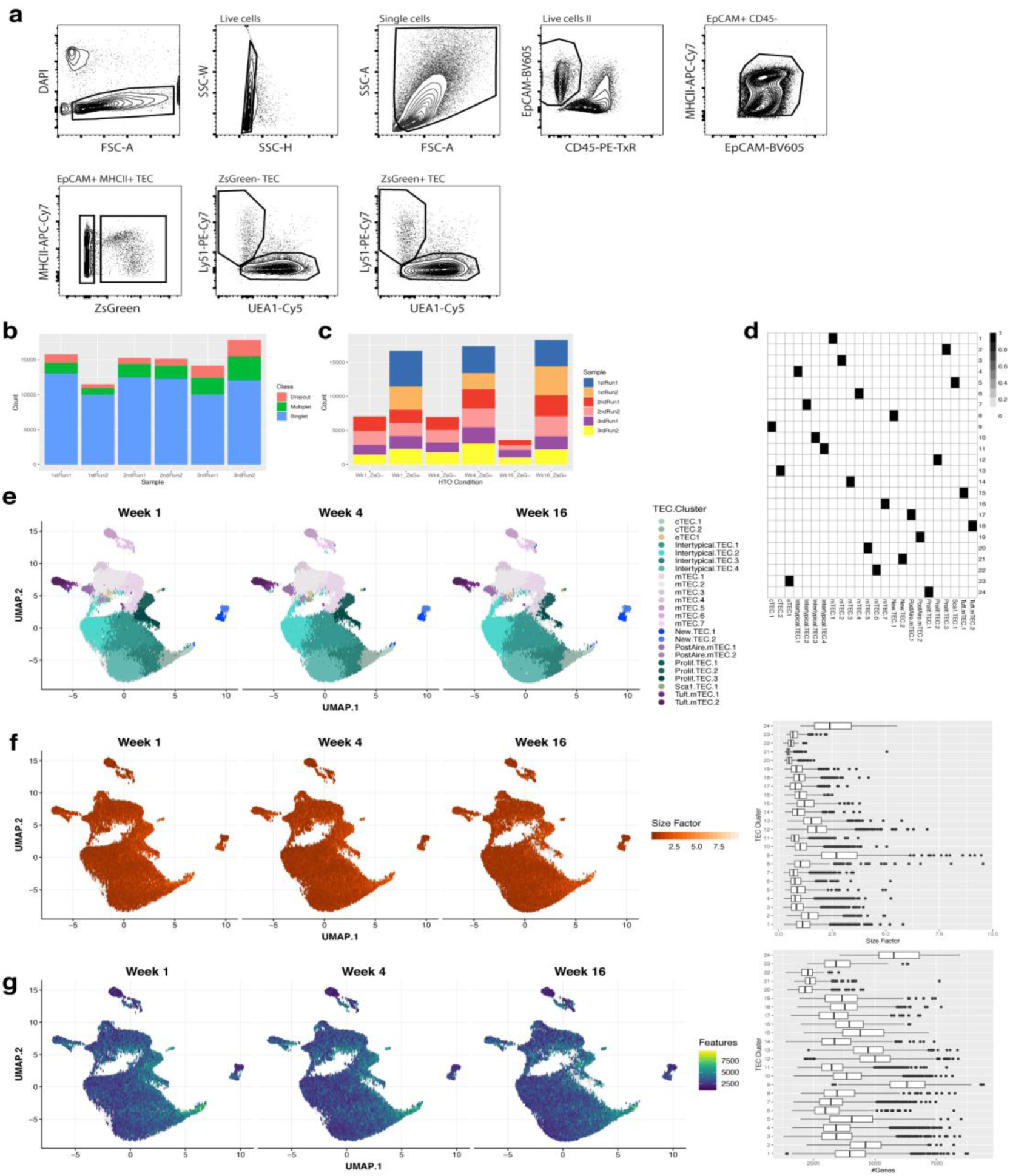
Quality control and summary of multiplexe d single-cell droplet RNA sequencing. (a) FAC-Sorting gating strategy for the isolation of ZsGreen +/− TEC subpopulations. (b) Multiplet detection using multiplexed hashtag oligos (HTO). Coloured bars denotes the number of cells in each sample (Chromium chip well), where either no (Dropout), multiple (Multiplet) or a single HTO was detected in a droplet. (c) The distribution of singlet cells across samples and experimental conditions (age and ZsGreen fraction). (d) A mapping of single-cell clusters onto equivalent ageing clusters. (e) Uniform manifold approximation and projection (UMAP). Points are single cells coloured by the assigned TEC subtype. Cells are split into panels based on the age of the mouse at the time of doxycycline treatment. (f) A UMAP split by mouse age showing the estimated deconvolution size factors. The boxplot on the right shows the distribution of size factors across single-cell clusters. (g) The number of detected genes (log expression > 0) in single cells overlaid on a UMAP and split by age. The boxplots on the right-hand side show the distributions of the number of detected genes for each single cell cluster.

**Supplementary Figure 9:**
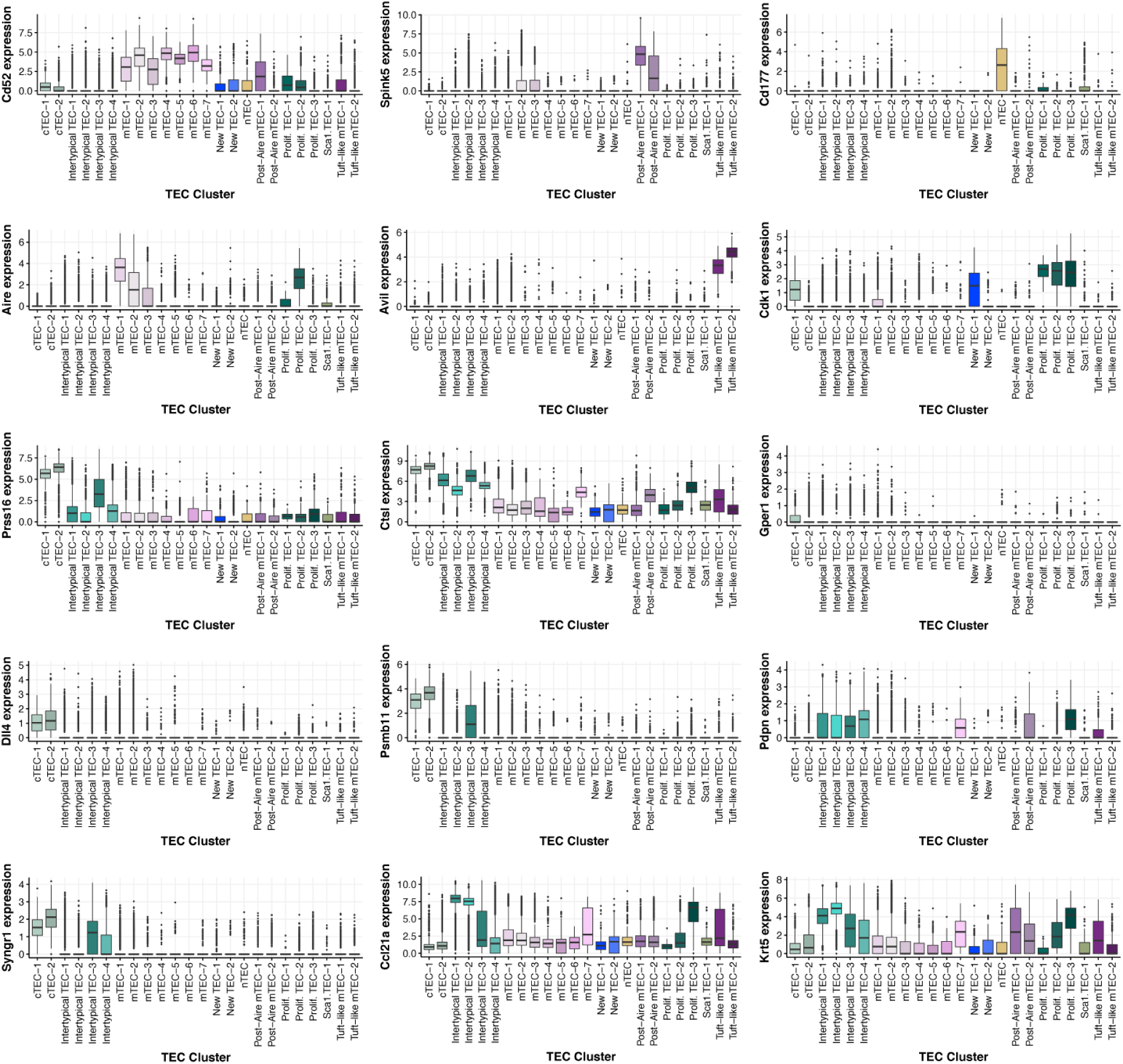
Marker gene expression profiles across TEC clusters from β-5t lineage-traced single cells. Boxplots showing the distribution of marker gene expression (y-axis) for TEC subtypes across TEC clusters (x-axis). Boxes are coloured by the inferred TEC subtype to which they belong.

**Supplementary Figure 10:**
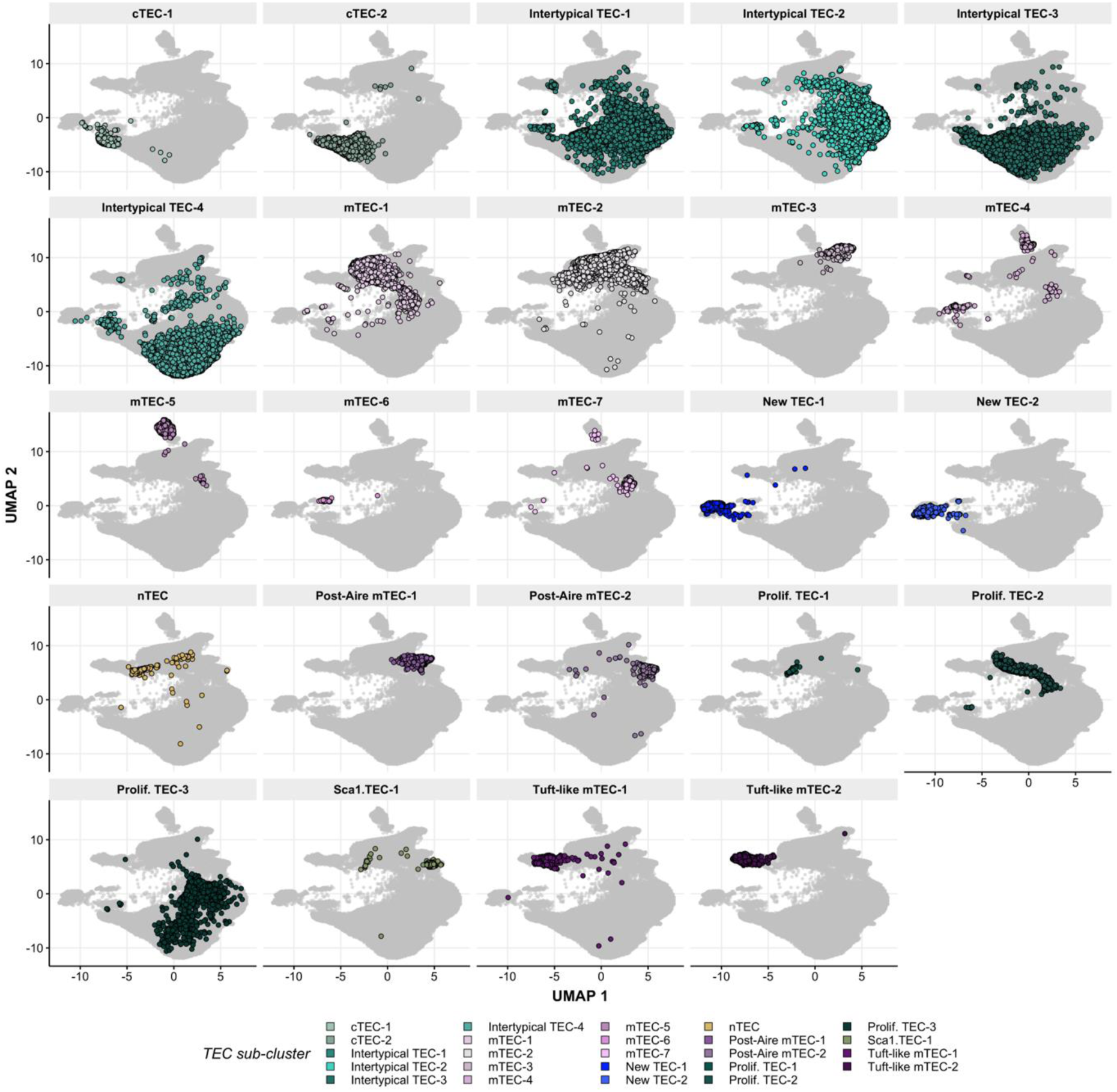
UMAP visualisation of TEC sub-clusters across all single-cells from lineage-traced thymi. Each panel is coloured according to the TEC subtype annotation and corresponds to Figure 4a.

**Supplementary Figure 11:**
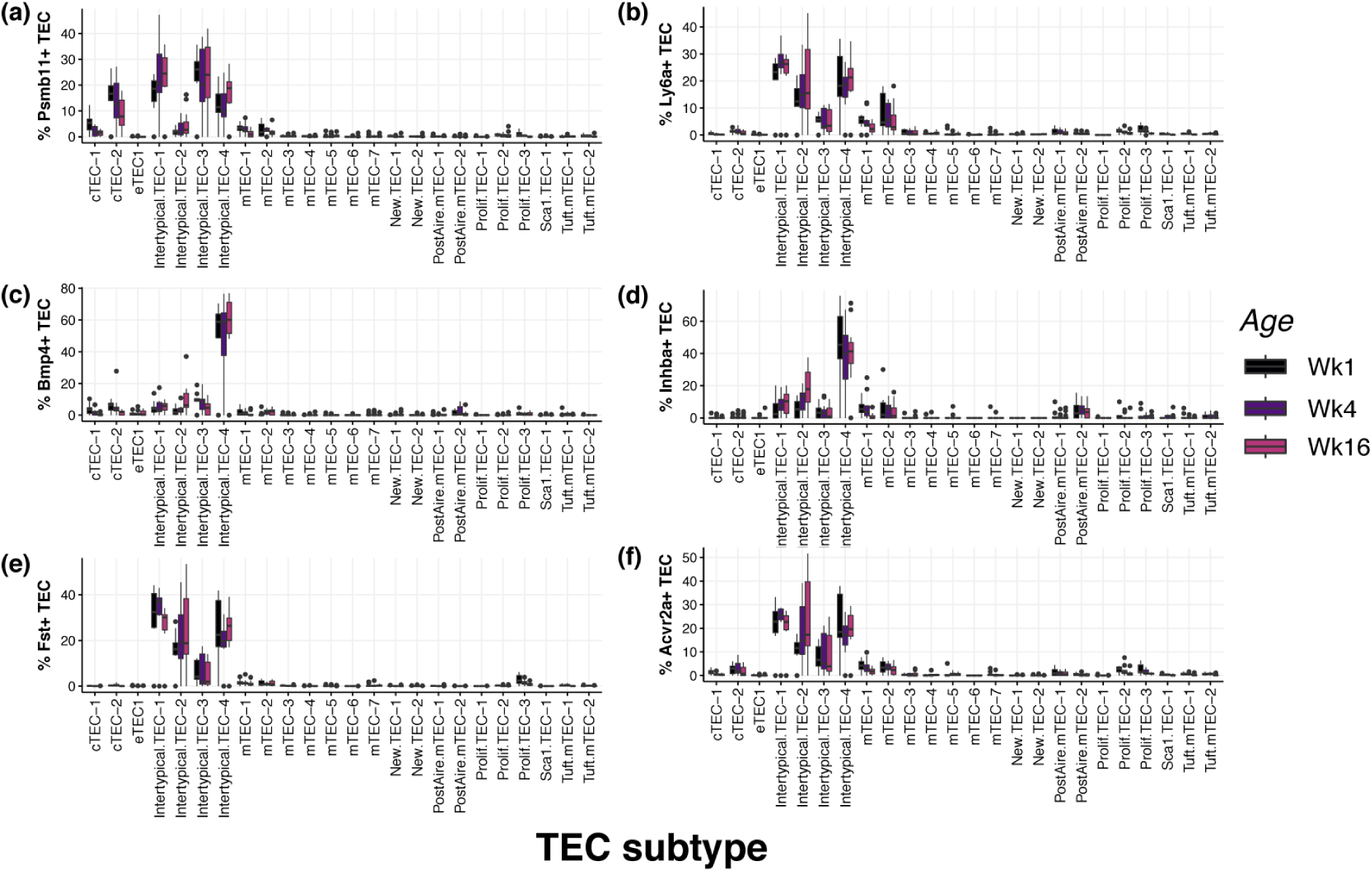
Boxplot showing the proportion of TEC in each subtype cluster which express key genes linked to thymic involution and TEC identity: (a) *Psmb11* (β5-t), (b) *Ly6a* (Sca1), (c) *Bmp4*, (d) *Inhba* (Activin A), (e) *Fst* (follistatin) and (f) *Acvr2a* (Activin A receptor 2a). Boxplots are coloured by age of dox-treatment administered to 3xtg^β5t^ mice.

**Supplementary Figure 12:**
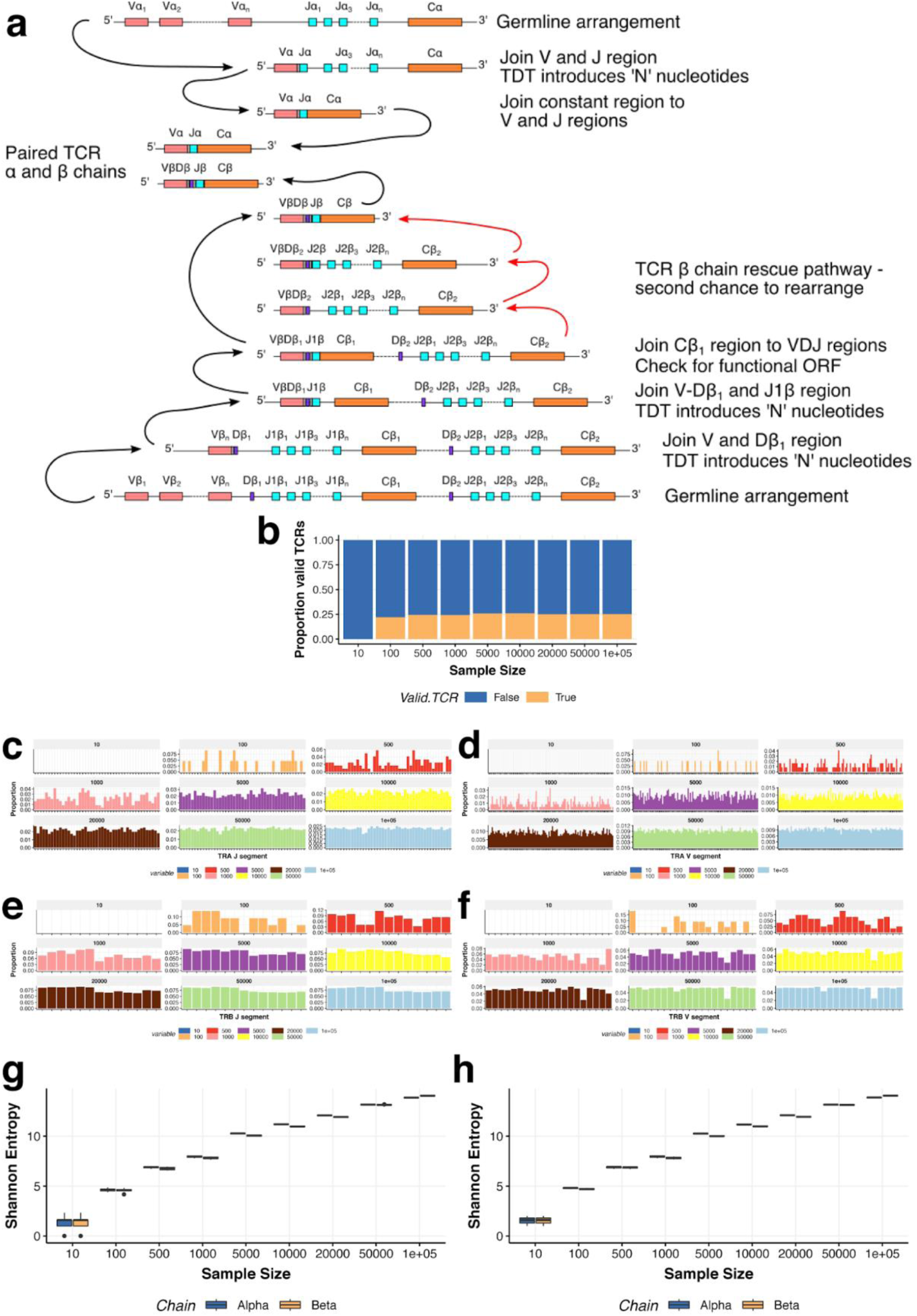
T cell receptor repertoire simulations. (a) A schematic of T cell receptor rearrangements used to design simulations. (b) Proportions of valid TCRs (y-axis) simulated at different sample sizes (x-axis). (c-f) Proportions of TCR alpha (c & d) and beta (e & f) chain segments from simulated TCRs at different sample sizes. (g-h) TCR repertoire diversity defined as the Shannon entropy across clonotypes in each of 10 (g) and 3 (h) independent TCR simulations. Entropy was calculated in each run using only the valid TCRs.

**Supplementary Figure 13:**
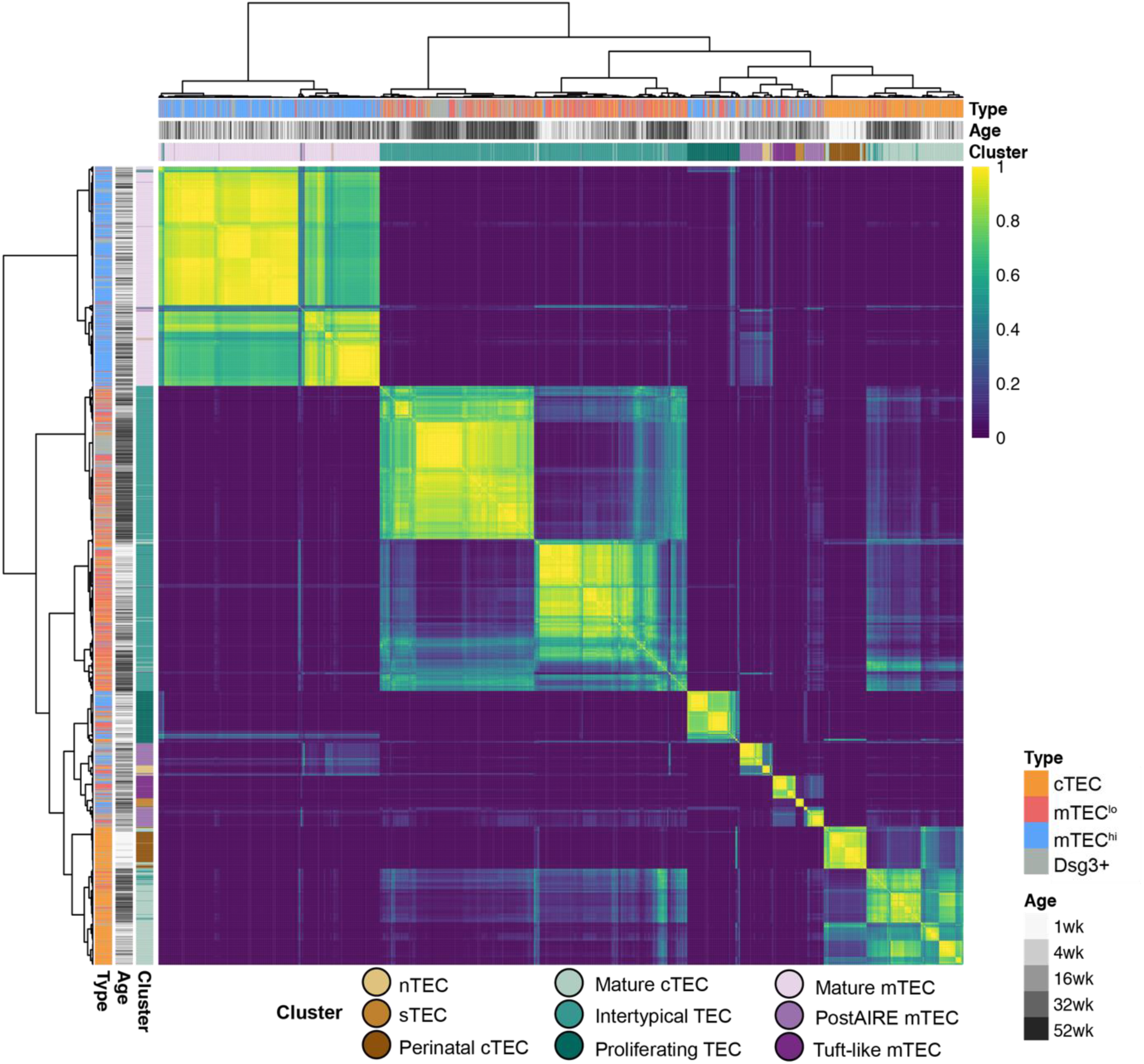
Consensus clustering of ageing single-cell TEC libraries. The heatmap shows the fraction of times that the libraries are co-clustered based on a variety of transformations, clustering methods and the number of features (Methods). The heatmap shows that the achieved clustering is robust to these different clustering methods.

